# Multivalent modulation of endothelial LRP1 induces fast neurovascular amyloid-β clearance and cognitive function improvement in Alzheimer’s disease models

**DOI:** 10.1101/2024.05.06.592767

**Authors:** Junyang Chen, Pan Xiang, Aroa Duro-Castano, Huawei Cai, Bin Guo, Xiqin Liu, Yifan Yu, Su Lui, Kui Luo, Bowen Ke, Lorena Ruiz Perez, Xiawei Wei, Qiyong Gong, Xiaohe Tian, Giuseppe Battaglia

**Affiliations:** Functional and Molecular Imaging Key Laboratory of Sichuan Province, Huaxi MR Research Centre (HMRRC), Department of Radiology, West China Hospital of Sichuan University, Chengdu, 610000, China; Huaxi Xiamen Institute of Medical Research, West China Xiamen Hospital of Sichuan University, Xiamen, Fujian 361021, China; Institute for Bioengineering of Catalunya (IBEC), The Barcelona Institute of Science and Technology, Barcelona (Spain); Department of Chemistry, University College London (UCL), London, UK; The Xiamen Key Laboratory of Psychoradiology and Neuromodulation, Xiamen, China, 361000; Department of Applied Physics, University of Barcelona, Barcelona, Spain; Laboratory of Aging Research and Cancer Drug Target, State Key Laboratory of Biotherapy and Cancer Center and National Clinical Research Center for Geriatrics, West China Hospital, Sichuan University, Chengdu 610041, China; Research Unit of Psychoradiology, Chinese Academy of Medical Sciences, Chengdu, Sichuan, China; Catalan Institution for Research and Advanced Studies (ICREA), Barcelona, Spain; Curapath, Av. Benjamín Franklin, 19, 46980 Paterna, Valencia

## Abstract

The blood-brain barrier (BBB) is a highly selective permeability barrier that safeguards the central nervous system (CNS) from potentially harmful substances while regulating the transport of essential molecules. Its dysfunction is increasingly recognized as a pivotal factor in the pathogenesis of Alzheimer’s disease (AD), contributing to the accumulation of amyloid-β (Aβ) plaques. We propose an AD therapeutic strategy targeting the BBB low-density lipoprotein receptor-related protein 1 (LRP1). We show that a multivalent scaffold with LRP1-specific peptide modulates the Aβ transport at the BBB. We show via detailed experimentation on AD model mice that this intervention markedly reduces Aβ deposits and prevents cognitive decline. This study marks a new approach to drug design, combining multivalent targeting with the regulation of membrane trafficking through advanced molecular engineering. Crucially, the therapeutic effects emerge from the multivalent nature of our proposed system. Furthermore, our findings underscore the paramount significance of the BBB in AD pathogenesis, particularly emphasizing the critical role of LRP1-mediated Aβ clearance in mitigating disease progression.

## Introduction

Alzheimer’s disease (AD) accounts for almost 70% of dementia cases among various types. AD pathophysiology is characterized by an accumulation of small peptides, Amyloid-β (Aβ), into fibrils and plaques, followed by hyperphosphorylation, misfolding, and aggregation into neurofibrillary tangles of another protein, tau. Both aggregates induce strong inflammatory responses, synaptic dysfunction, and neuronal injury, causing considerable brain damage and impairing cognitive processes (1,2). Alongside these, the brain vasculature network, or blood-brain barrier (BBB), plays a critical role during AD progression and possibly initiation (3-6). The BBB consists of aligned endothelial cells supported by pericytes and astrocytes, forming the densest vascular network in the body, with about one capillary per neuron (3). The BBB poses a significant challenge in pharmacology, impeding the penetration of numerous drugs and complicating treatment discovery for neurological disorders (3). Most AD patients experience various vascular dysfunctions (7), which may be linked to Aβ (8) and tau (9-11) or occur independently of both (12). The low-density lipoprotein receptor-related protein 1 (LRP1) is possibly the most studied receptor for both Aβ (1, 13-15) and tau (16,17) processing. Endothelial LRP1 plays a vital role in removing Aβ, and its expression diminishes with age. This decrease is more pronounced in AD patients and animal models, where BBB LRP1 levels are almost undetectable (18-22). The down-regulation of LRP1 is strongly correlated with impairment of the BBB and cognitive decline (23-28).

Proper regulation of LRP1 levels in endothelial cells is crucial for preventing the progression of Alzheimer’s disease (AD). Despite this, the mechanisms that maintain appropriate LRP1 levels on the basolateral surface of endothelial cells remain unclear. Research has shown that Aβ/LRP1 complexes are trafficked by phosphatidylinositol-binding clathrin assembly protein (PICALM) and clathrin, processed within RAB5 and RAB11 compartments, and released into the bloodstream via transcytosis (24-31). We have observed (32,33) that LRP1 receptors also cross the blood-brain barrier (BBB) through a Rab5/PICALM-independent pathway, involving sequential collective endocytosis, tubular vesicle formation stabilized by PACSIN2 (Syndapin-2), and exocytosis (schematized in Fig1a). Our findings demonstrate that synthetic nanoparticles with varying LRP1 ligand numbers (32) or Aβ structures with different aggregate sizes (33) dictate LRP1 transport via Rab5 or PACSIN2. High-avidity cargo directs LRP1 into Rab5-endosomes, down-regulating LRP1, while mid-avidity cargo directs LRP1 into PACSIN2 tubular vesicles, up-regulating LRP1 by avoiding endosomes and acidification (32,34).

**Fig. 1.**
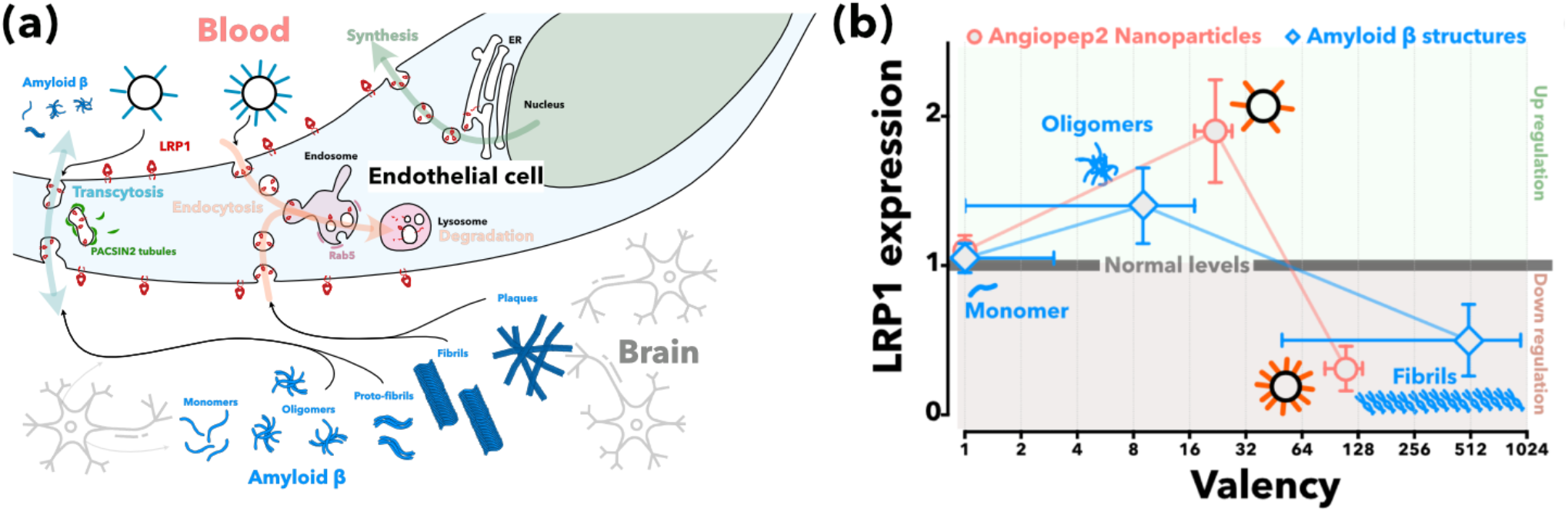
Schematics of the LRP1 shuttling across brain endothelial cells (**a**) following either a PACSIN2 or Rab5 pathway and its relation with multivalent cargo. LPR1 expression in brain endothelial cells (**b**) as a function of the cargo valency. This latter represent the average number of the ligands per cargo that interact with LRP1, with n=1 representing the single peptide. The data are replotted from refs 32 and 33.

This division indicates that as Alzheimer’s disease progresses and individuals age, larger Aβ structures with stronger binding to LRP1 reduce LRP1 levels, leading to impaired Aβ clearance. On the other hand, using mid-avidity cargo to target LRP1 can increase LRP1 levels, counteracting the impact of Aβ on the BBB’s equilibrium. Therefore, we propose repurposing the multivalent carrier with LRP1-binding Angiopep2 peptide-functionalized on P[(OEG)10MA]20-PDPA120 polymersomes (A39-POs) optimized for ‘super-selective’ BBB targeting and PACSIN2-mediated transcytosis (32-35) to promote LRP1 up-regulation and enhance Aβ clearance, potentially serving as a disease-modifying treatment.

## Results and discussion

To fully understand the role of LPR1 in mediating Aβ transport, we followed its expression alongside other markers and Aβ in both AD model APP/PSEN1 and wild-type animals. We employed the Enzyme-Linked Immunosorbent Assay (ELISA) and imaging methods (**Fig. S1**) to conduct a comparative analysis of the whole brain before and after its fractionation in vasculature and parenchyma. We measured Aβ levels in AD model mice against wild-type controls over a lifespan from 3 to 12 months. We also measured key proteins involved in receptor-mediated transcytosis and endocytosis, tight junction proteins, neurons, and other associated proteins. We discerned the temporal changes of these proteins from a macroscopic whole brain and BBB. **Fig. 1a** and **Fig. 1b** display the aggregate Aβ levels in the brain, showing a marked rise in Aβ in the AD models with age, a trend particularly pronounced between 6 to 12 months. A measurable disparity in BBB Aβ levels between the AD model mice and the wild-type controls is evident in Fig. 1c, revealing significantly higher Aβ in the brain’s vasculature of wild-type mice, with notable differences emerging at all lifespan stages, emphasizing the pathological hallmarks of AD with its deficient BBB transport capacity.

**Fig. 1.**
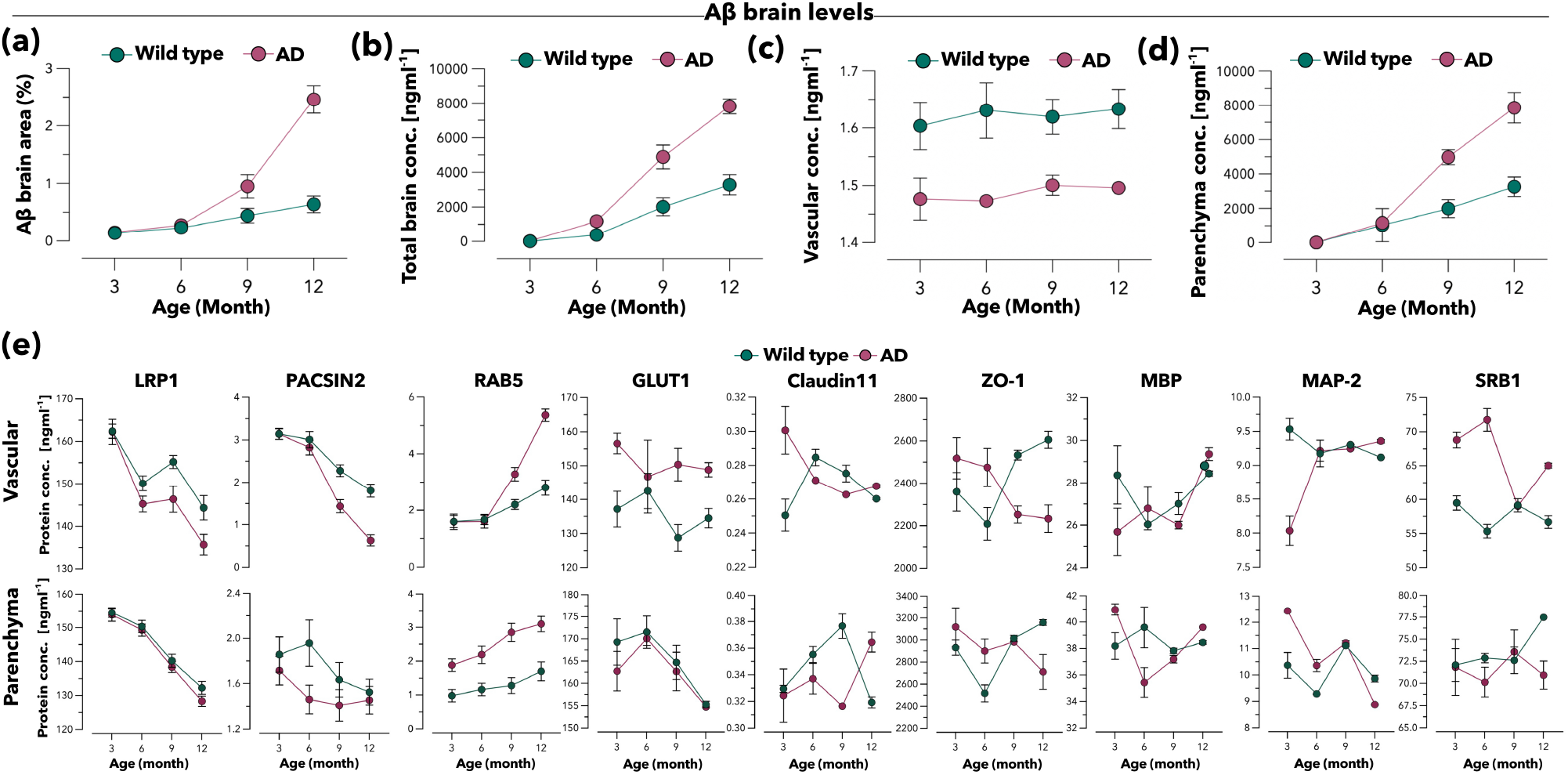
Comparative analysis of Aβ accumulation and protein expression profiles in AD and wild-type mice. Quantitative representation of Aβ-positive areas (**a**) in brain coronal sections, identified through immune histochemistry (IHC), in both AD model and wild-type mice. Quantitative measurement of Aβ concentrations via ELISA in both AD and wild type mice of the whole-brain (**b**), its vasculature (**c**) and parenchyma (**d**). ELISA quantification of both brain vasculature and parenchyma (**e**) in AD and wild type mice of key proteins involved in trafficking (LRP1, PACSIN2, RAB5), metabolism (GLUT1), tight junction proteins (ZO-1, CLDN11), and other stress-related proteins (MBP, MAP-2, SRB1). For all analyzes (n ≥ 3)

Conversely, **Fig. 1d** delineates the parenchymal Aβ concentration, indicating a significant surge in the AD models, especially between 9 to 12 months. The following ELISA assessment shows the changes in various protein levels in the brain’s vascular and parenchymal regions in AD model mice and wild-type controls over the same period. We focused on LRP1, PACSIN2, RAB5, GLUT1, Claudin 11, ZO-1, MBP, MAP-2, and SRB1, which are essential for processes like transcytotic transport, lipid transport, glucose metabolism, and the maintenance of cellular structure and integrity. Notable variations were observed throughout the months, with proteins like MBP and MAP-2, essential for neural integrity, displaying divergent trends potentially indicative of neurodegenerative alterations characteristic of AD (**Fig. 1e**). We also observed that the transcytosis-related protein s LRP1 and PACSIN2 showed marked reductions on vascular endothelial cells as AD advanced. Meanwhile, RAB5, integral to endocytosis, and GLUT1, essential for glucose transport across the BBB, demonstrated noticeable increments correlating with AD progression. Fluorescent images showed the same trend (**Fig. S2**).

Following the clue upon analyzing the brain vascular protein variation, we then use confocal microscopy images that showcase the spatial localization of LRP1 and A β at the BBB endothelium (CD31) and pericytes (CD146) in 3-month-old and 12-month-old AD brain samples. **Fig. 2a** (3-month-old), Aβ (red) is highly co-localized with LRP1 (white) on the surface of blood vessel cells of the BBB (green), suggesting active involvement of LRP1 in Aβ transport and clearance at a younger age, with less Aβ accumulation around the vessels.

**Fig. 2.**
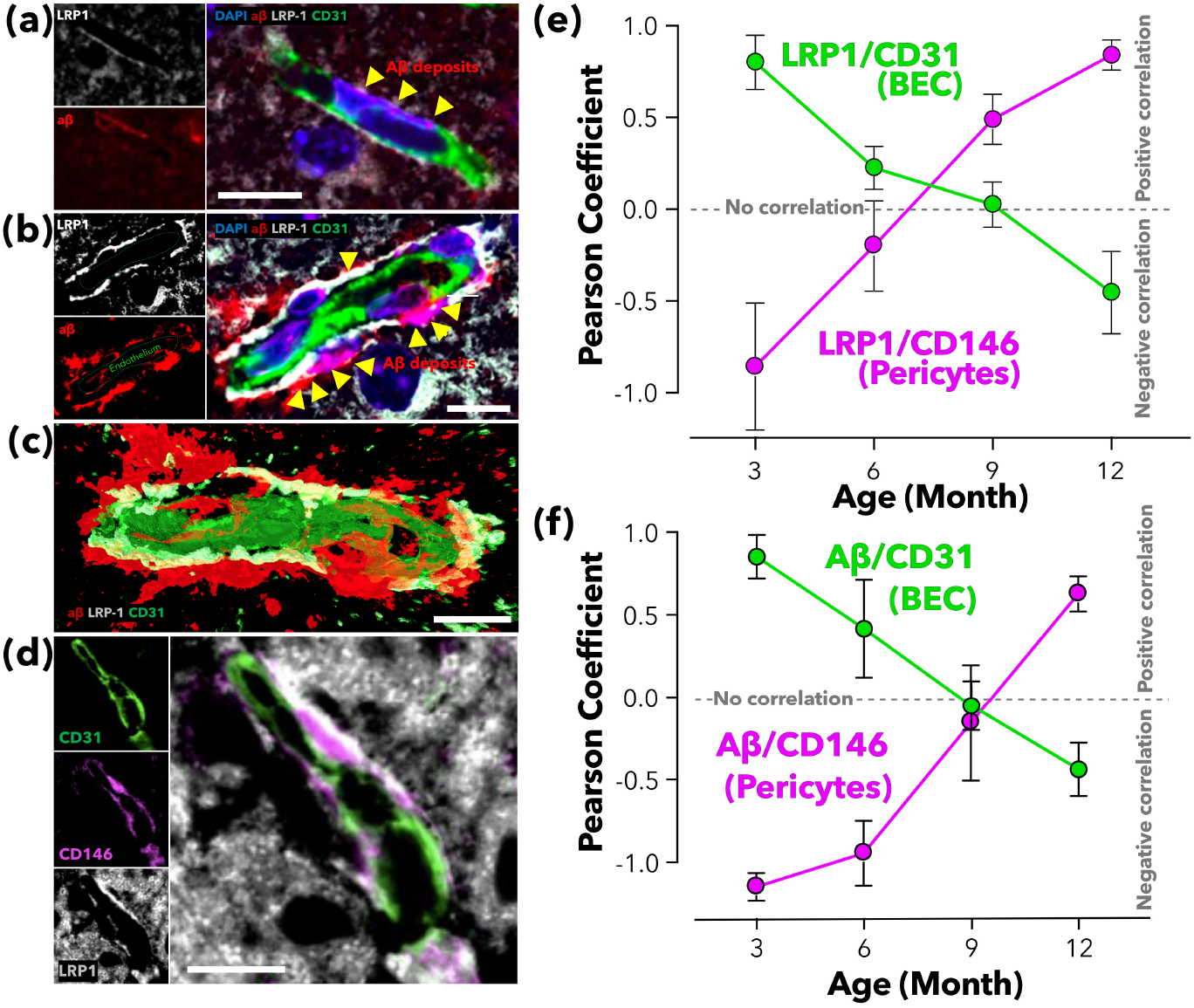
Confocal microscopy and co-localization analyzis of brain endothelial cells, pericytes, and Aβ in AD mice of different ages. LRP1 (grey), Aβ (red), and BBB endothelium (CD31, green) in 3-month-old (**a**) and in 12-month-old (**b**) AD mice brain samples. 3D rendering (**c**) of the 12-month-old AD brain section highlighting the spatial relationship between the different markers. In later AD stages, increased Aβ accumulation near the BBB and reduced co-localization of CD31 with LRP1 are observed. CD31, CD146, and LRP1 in a 12-month-old AD brain sample (**d**). Scale bar = 10 μm. Pearson correlation coefficients over different ages for LRP1 (**e**) and Aβ (**f**) with endothelial cells (CD31) and pericytes (CD146), showing changing interaction patterns with ageing.

**Fig. 2b** shows 12-month-old brains, and its corresponding 3D image (Fig. 2c) indicates a noticeable increase in Aβ deposition on the basal side of the BBB vessels. The co-localization of LRP1 with Aβ appears to decrease, potentially signifying impaired LRP1-mediated clearance of Aβ as AD progresses. At later stages of AD, the increased Aβ accumulation and reduced association with LRP1 may affect BBB function and promote disease pathology. Interestingly, further image (**Fig. 2d**) suggests LRP1 appears predominantly deposited around the pericytes on the exterior side of the BBB blood vessels. We further analyze representative brain sections from 3-12 months, labeling LRP1, Aβ, and pericytes and endothelium markers CD146 and CD31, respectively. We performed a co-localization analysis and calculated the Pearson correlation coefficient between LRP1 and CD31, and between LRP1 and CD146. In **Fig. 2e**, the results show a trend where the association between LRP1 and endothelial cells weakens over time (green line), while its correlation with pericytes appears to strengthen (purple line). Similarly, the same analysis with Aβ shows a Pearson correlation coefficient over time with both endothelial cells (green line) and pericytes (purple line) an inverse association (**Fig. 2f**). Combining the above ELISA and confocal evaluation, the shift in LRP1 from BBB vascular endothelium to pericytes with aging underscores a potentially pivotal role in the pathophysiology of AD. This progression suggests a decrease in LRP1-mediated Aβ clearance at the BBB endothelial level, with a concomitant increase associated with pericytes, which may impact AD progression.

Our previous studies (32, 34, 36) found that the peptide Angiopep2 can target LRP1. We also discovered that the efficiency of crossing the BBB is higher when the ligand is present on a multivalent scaffold. Our research has shown that the transportation of LRP1 across the BBB occurs through transcytosis (32). This process involves collective endocytosis and exocytosis and is regulated by a BAR domain protein called PACSIN2. Our observation suggests that cargo with an intermediate number of ligands works best with this transportation mechanism. This includes synthetic cargos like A_39_-PO polymersomes (14) and natural cargos like Aβ assemblies (15). Both multivalent units trigger PACSIN2-mediated transcytosis. In both synthetic cargo and Aβ assemblies, PACSIN2-mediated transcytosis coincided with a temporary up-regulation of the LRP1 receptor (14, 15). Therefore, we hypothesize that using LRP1 multivalent targeted nanoparticles may restore the function of LRP1 in transporting Aβ out from the brain and potentially have the effect of clearing Aβ deposits in the AD model. We prepared and characterized Angiopep2-P[(OEG)_10_MA]_20_-PDPA_120_ polymersomes bearing 39 ligands per particle from now on, referred to as A_39_-POs (**Fig. S3**). APP/PS1 transgenic AD mouse was intravenously (i.v.) injected with A_39_-POs (10 g/L, 200 μL), and ELISA analyzed the whole brain. The Aβ contents showed a 40% reduction after a short administration period (**Fig. 3a**), from 7500 ng/mL to 4500 ng/mL. A parallel immune histochemistry (IHC) analyzis confirmed that the Aβ area fraction also decreased (**Fig. 3b** and **Fig. S4**). The brain reduction is mirrored by an increase in blood Aβ concentration, proving that the Aβ is shuttling from the brain to blood (**Fig. 3c**). Furthermore, we employed Positron Emission Tomography-Computed Tomography (PET-CT) to assess the clearance of Aβ in the brains of live animals. We utilized (E)-4-(2-(6-(2-(2-(2-^18^F-fluoroethoxy)ethoxy)ethoxy)pyridin-3-yl)vinyl)-N-methyl benzamine ([^18^F]AV-45), which has been previously demonstrated to strongly correlate with the presence and density of Aβ (22). We thus imaged the brains of young (3-month-old) wild-type mice as controls. The PET-CT scans revealed that the brain of a 12-month-old APP/PS1 mouse exhibits an intense Aβ signal. In contrast, this signal sharply decreased after the treatment of A_39_-POs (**Fig. 3d**). Following the administration of A_39_-POs, the Standardized Uptake Value (SUV) associated with Aβ reduction was 46.25% (overnight) (**Fig. 3e**). We finally performed tissue clearing on brains of Sham APP/ PS1 12-month-old and A_39_-POs treated 12-month-old APP/PS1 mice. The Aβ (red) and BBB (green) of these brains were labeled, respectively (**Fig. 3e**). Brains of mice treated by A_39_-POs had less Aβ signal relative to the Sham APP/PS1 brain. 3D brain images were nested into the Amira software companion brain partitioning model and paired (**Fig. 3g**). Aβ volume in 14 brain regions of the mouse brain was measured separately (**Fig. 3h**). A total of 41% reduction in mice brain Aβ volume after treatment with A_39_-POs. Lastly, coronal Aβ distribution is shown as a heat map in **Fig. 3i**.

**Fig. 3.**
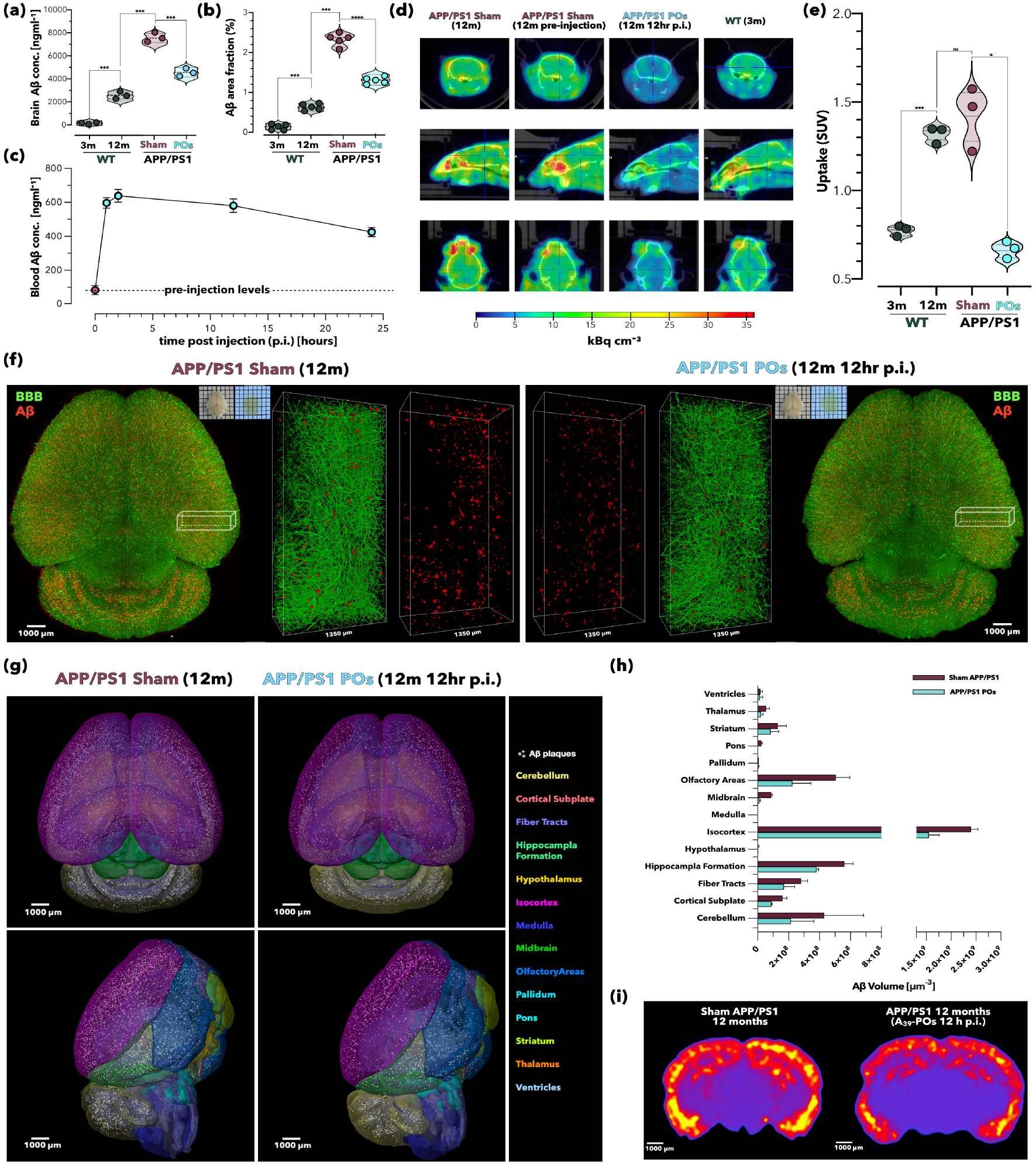
A_39_-POs reduce Aβ levels in the brains of AD mice. ELISA measurement of whole-brain Aβ levels 2 hr post A_39_-POs administration (**a**) revealed significant reductions, from ∼7500 ng/ml to ∼4500 ng/ml. Quantification of Aβ positive area fraction on IHC stained coronal brain sections (**b**). Temporal Aβ concentration profile in blood (**c**) post A_39_-POs injection, sampled at 0, 1, 2, 12, and 24 hr. PET-CT whole body imaging (**d**) of wild type (3 and 12 months old) and APP/PS1 (12-month-old pre- and 12 hr post A_39_-POs injection) mice following [^18^F]AV-45 (2.8-3.2 MBq) administration 1 hr prior to scanning. Standardized uptake value (SUV) of [^18^F]-AV45 (**e**) indicating a significant post treatment Aβ level decrease in the brain. 3D rendering from light sheet fluorescence imaging (**f**) of brain after tissue clearing shows reduced Aβ signals (in red) in A_39_-POs treated mice at 12 hr. Brain region mapping (**g**) with noted reduction in Aβ plaques in the isocortex. Brain segmentation into 14 regions revealing predominant Aβ distribution in the isocortex, with a total volumetric reduction of 41% post A_39_-POs treatment (**h**). Thermal map representation of Aβ fluorescence intensity across a single coronal brain layer (**i**). Tissue clearing experiments were conducted at least twice per group. Statistical significance was determined using One-way Analyzes of Variance (ANOVA), *p<0.05, **p<0.01, ***p<0.001, ****p<0.0001, n ≥ 3.

The above outcomes motivated us to re-study the BBB vasculature phenotype after A_39_-POs treatment. We first observed a reserve of the co-localization of LRP1 with CD31 in the mouse brain, as shown in **Fig 4a** and **Fig. S6**; the overlap of LRP1 and BBB endothelial cells (CD31) returned to the state of wild type. We thus analyzed the Aβ content in brain vascular endothelial cells and parenchyma and found a significant increase in its content in BBB and a notable decrease in parenchyma (**Fig. 4b**). The ELISA assessments again are performed to indicate proteins in both vascular and parenchymal after A_39_-POs treatment in 12-month-old AD mice (**Fig. 4c**). As discussed previously, the analysis focuses on the concentrations of various proteins such as LRP1, PACSIN2, RAB5, GLUT1, Claudin 11, ZO-1, MBP, MAP-2, and SRB1. The nanomedicine drug cleared Aβ and caused a rapid change in the BBB phenotype by up-regulating PACSIN2 and down-regulating RAB5. This finding is consistent with our confocal data, which shows that LRP1 relocates to blood vessels. The morphology of LRP1 (white), observed under a stimulated emission depletion (STED) microscope, showed clustered distribution upon the vessel wall (green), suggesting robust ongoing transcytosis. We also observed that the Aβ deposition (red) around the BBB disappeared, and confocal images displayed a large amount of Aβ signal in the vascular lumen (**Fig. 4d**).

**Fig. 4.**
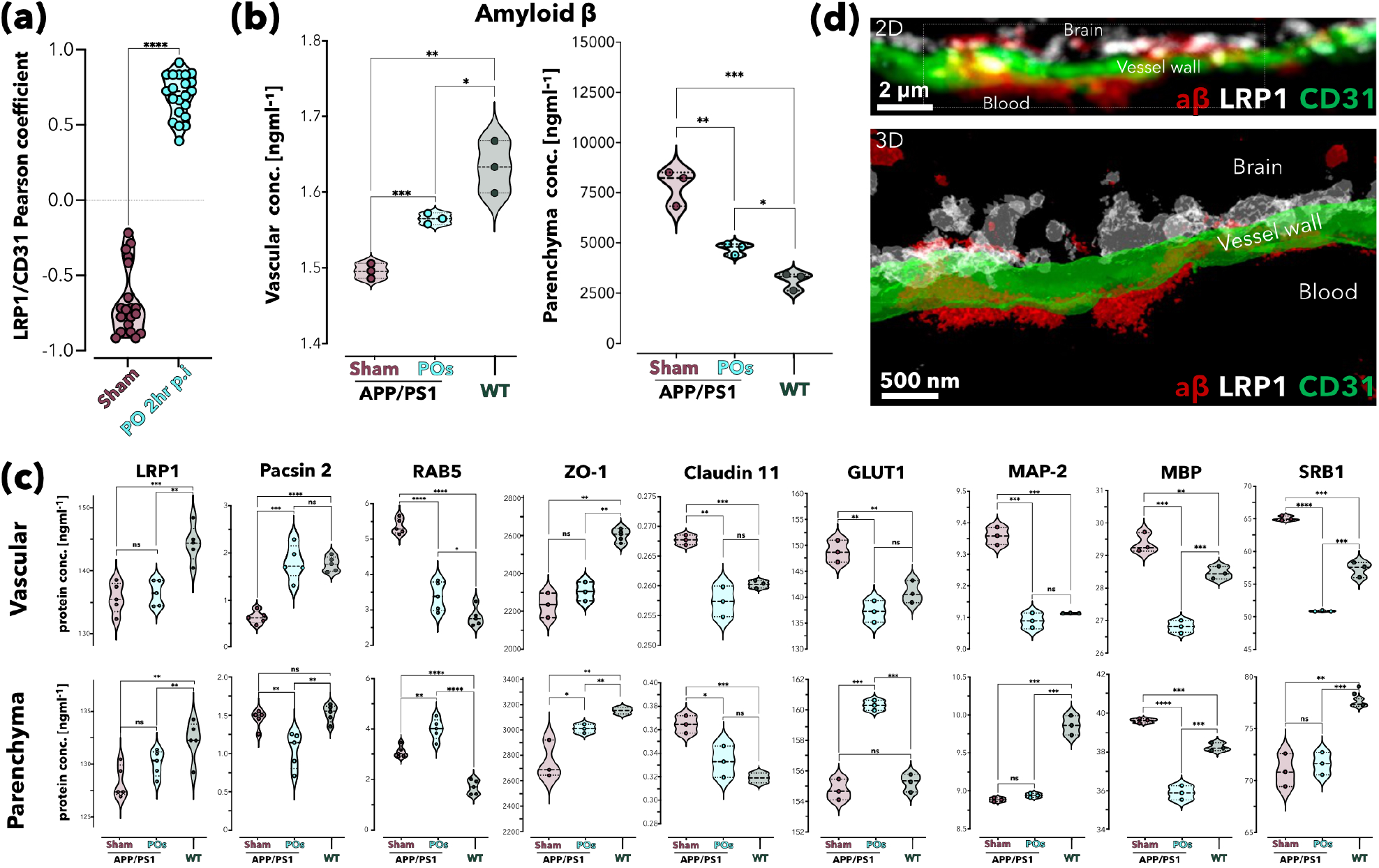
A_39_-POs treatment restores healthy BBB phenotype. Pearson correlation coefficients for the colocalization of LRP1 and BBB endothelial cells (CD31) pre- and post A_39_-POs administration (**a**). Analysis performed on images derived from three independent experiments, with 6-7 vessels studied per trial. Quantification of Aβ content in cerebrovascular endothelial cells and brain parenchyma (**b**) (n = 3, statistical analysis performed using unpaired t-tests, **** p<0.0001). ELISA measurements (**c**) of vascular and parenchymal proteins in 12-month-old wild type, Sham APP/PS1, and APP/PS1 post A_39_-POs treatment mice, encompassing LRP1, PACSIN2, RAB5, GLUT1, Claudin 11, ZO-1, MBP, MAP-2, and SRB1. STED microscopy (**d**) of LRP1 (white) on the vessel wall (green), indicative of active transcytosis. Post-treatment, Aβ deposits around the BBB (red) are cleared with notable Aβ signal presence within the vascular lumen. For (b) and (c), statistical significance determined using one-way ANOVA, *p<0.05, **p<0.01, ***p<0.001, ****p<0.0001, n ≥ 3.

Finally, we showed the effect of A_39_-POs administration on animal cognition using enhanced spatial learning and memory. As indicated in **Fig. 5a**, the stage I data showed that compared to the Sham APP/PS1 group, the APP/PS1 POs group could find the escape platform via a shorter path, indicating that the mice have a similar search strategy to the wild types. Their spatial learning and memory abilities are superior to those of the Sham APP/PS1 group. The escape latency (time taken to find the escape platform) of the APP/PS1 POs group was also shorter. As the experiment days increased, the time animals took to find the platform gradually decreased, suggesting that they made progress in learning and remembering the platform’s location. A shorter escape latency indicates the animals’ better spatial learning and memory abilities. Although the relationship between swimming speed and spatial learning and memory abilities is weak, analyzing swimming speed can rule out the impact of the animals’ motor abilities or fear during the experiments. For all parts, there was no statistical difference in the swimming speeds of the three groups of mice.

**Fig. 5.**
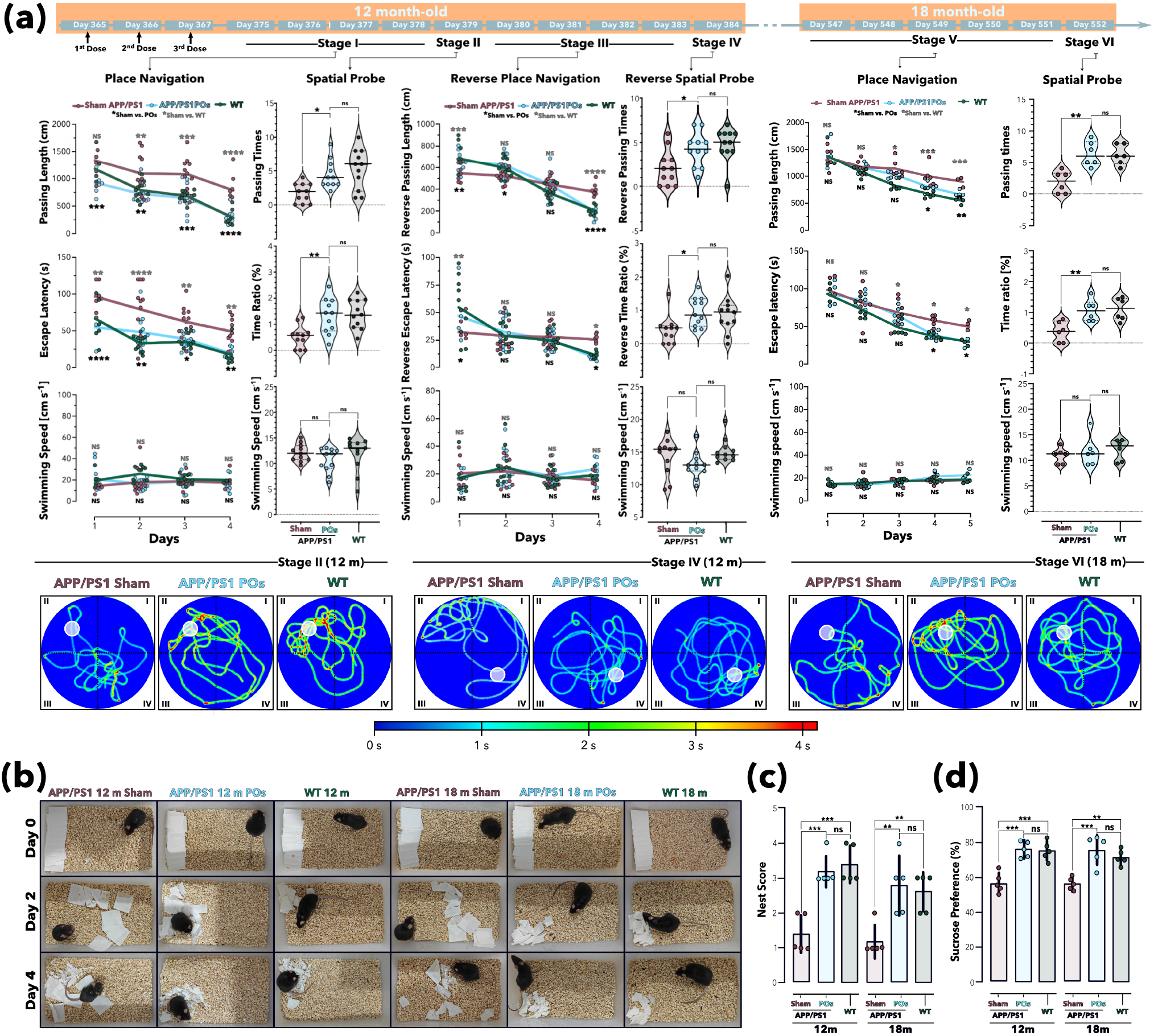
Behaviour tests demonstrated that A_39_-POs treatment improved performance of AD mice. Morris Water Maze test **(a)** which mice injected with saline (Sham APP/PS1 group and WT group, 200 μL) or A_39_-POs (APP/PS1 POs group, 10 g/L 200 μL) once daily for the 365th - 367th mornings of lifespan. Recovery was execute for 1 week under original rearing conditions. Place navigation (Stage I) test occurred on days 375th - 378th, showing a gradual decrease in escape length and escape latency for finding the platform in all groups, with the APP/PS1 POs group matching the WT level and significantly outperforming the Sham APP/PS1 mice. During the spatial probe (Stage II), both the APP/PS1 POs and WT groups demonstrated more passing times and a higher percentage of time spent at the escape platform’s original location. The reverse place navigation (Stage III) trial from days 380th - 383th, with the platform moved to the opposite side (IV quadrant), saw the APP/PS1 POs and WT groups initially taking longer, indicating stronger spatial memory from stage I and stage II. However, their reverse passing length and escape latency decreased rapidly over time and were significantly lower than those of the Sham APP/PS1 mice. On day 384, in the reverse spatial probe (Stage IV) test without the platform, A_39_-POs treated mice still outperformed the Sham APP/PS1 group. After 180 days, mice were re-performed for place navigation and spatial probe (Stage V and Stage VI). Conditions were consistent with stage I and stage II. Six months after the injection of A39-POs, the mice could still find the escape platform with shorter time in the place navigation experiment. They stay longer at the correct location for and traverse the original platform location more times in the spatial probe. The performance of mice injected with A39-POs (6 months p.i.) was close to that of the wild type, and both were better than that of the Sham APP/PS1 group (a). Place navigation trials (Stages I, III and V) were analyzed using two-way ANOVA, while spatial probe trials (Stages II, IV and VI) comparisons used one-way ANOVA. Significance levels are denoted as *p<0.05, **p<0.01, ***p<0.001, ****p<0.0001, with n ≥ 11 (for all 12-month-old mice) and n ≥ 6 (for all 18-month-old mice). Nest-construction images (**b**), nest score (**c**) and sucrose preference (**d**) were record for APP/PS1 Sham, APP/PS1 POs and wild type group at two time post injection.

When the experiment entered stage II, and the escape platform was removed, the APP/PS1 POs group crossed the platform more times and spent a significantly larger percentage of time at the platform’s original location than the Sham APP/PS1 group, reflecting stronger long-term memory of the platform’s location. When the escape platform was placed on the opposite side of its original location. Initially, longer search times reflected a long-term memory of the original platform’s location. Still, as the number of training sessions increased, the group with stronger learning abilities (APP/PS1 POs group) showed a higher rate of reduction in path length and escape latency. In the last two days of this stage, animals treated with A_39_-POs showed shorter escape paths and shorter escape latencies. When the escape platform was removed, the APP/ PS1 POs group stayed longer at the escape location. It crossed this location more times, reflecting the stronger memory abilities of the animals treated with A_39_-POs, with nearly no difference in swimming speeds during this stage. Six months after the mice were treated, we performed a water maze experiment on these mice to evaluate the persistence of cognitive improvement in mice treated with A_39_-POs. Place navigation and spatial probe tests (Stage V and Stage VI) were performed on the mice using the same methods and conditions as in Stage I and Stage II. Between-group comparisons showed that the cognitive enhancement provided by A_39_-POs persisted in APP/PS1 mice, and A_39_-POs treated mice demonstrated a level of cognitive improvement similar to that of wild type, which was significantly promoted compared with sham APP/PS1.

Enhancing the quality of life is a crucial objective in AD treatment and improvement. To assess the quality of life in mice, we conducted nest construction (**Fig. 5b** and) and sucrose preference experiments following the Morris water maze test (Stage IV and Stage VI). Nest-construction behavior is commonly used to evaluate daily activities, fine motor skills, cognition, and emotional state in mice with cognitive impairments. For mice, a high-quality nest provides insulation and protection against enemies, serving as an important place for them to feel secure, thus reflecting their executive function level as well. The sucrose preference experiment measures animals’ hedonic response towards sweetness by providing pure water and low-concentration sucrose solution to indicate pleasure deficit symptoms in rodents. After A_39_-POs treatment, APP/PS1 mice exhibited comparable levels of nest-building ability (nest score in **Fig. 5c**) and happiness perception ability (sucrose preference in **Fig. 5d**) to wild-type levels, immediately after treatment as well as six months post-injection, which is significantly higher than those observed in sham APP/PS1 group.

Overall, the behavioral experiments indicate that animals treated with A_39_-POs showed improved navigation and learning capabilities, exhibited cognitive enhancements, and gained a higher quality of life attributed to the rapid clearance of Aβ following A_39_-POs treatment.

## Conclusion

Our study represents a substantial advancement in AD therapeutics by applying a multivalent targeting approach, specifically utilizing LRP1-targeting peptides functionalized polymersomes to modulate its membrane trafficking. This strategy has demonstrated remarkable efficacy in achieving rapid Aβ clearance, precipitating an immediate and beneficial alteration in the BBB phenotype and resulting in a pronounced reversal of AD pathology. The therapeutic intervention led to a significant reduction in Aβ deposits, which was accompanied by a swift modification in both vascular and parenchymal protein profiles. These changes are indicative of a restored BBB function, underscoring the pivotal role of the BBB in AD pathogenesis and the potential of LRP1-mediated Aβ clearance as a therapeutic target. The cognitive improvements observed in AD model mice further validate the effectiveness of our approach, highlighting its capacity to mitigate pathological hallmarks and enhance neurological function. The strategic combination of targeted Aβ clearance with the modulation of BBB phenotype characterizes this promising new direction for AD treatment. By integrating advanced molecular engineering techniques with multivalent targeting, our approach leverages the multivalent nature of LRP1-functionalized polymersomes to achieve superior therapeutic outcomes. The rapid and effective clearance of Aβ, coupled with the restoration of BBB integrity, presents a compelling case for this strategy as a foundational framework for future AD therapies.

In conclusion, our study underscores the transformative potential of multivalent targeting and BBB modulation in treating AD. The deployment of LRP1-targeting polymersomes has facilitated rapid Aβ clearance and initiated significant changes in the BBB, culminating in improved cognitive outcomes. This innovative therapeutic paradigm offers a promising pathway for developing effective clinical interventions, addressing vascular contributions to AD, and ultimately enhancing patient outcomes.

## Acknowledgments

This study was supported by the National Key R&D Program of China (2022YFC2009900), the Alzheimer’s Association New to the Field award, ERC Consolidator grant H2020-ERC-2018-CoG (769798 CheSSTag), Plan de Recuperacion Nacional Biotech for Health Project (ADNano), Activitat científica dels grups de recerca de Catalunya (SGR-Cat 2021), and Spanish Research Agency Proyectos I+D+I PID2020-119914RB-I00.

## References

1. H. Hampel et al., The Amyloid-beta Pathway in Alzheimer’s Disease. Mol Psychiatry 26, 5481–5503 (2021).

2. E. Karran, B. De Strooper, The amyloid hypothesis in Alzheimer disease: new insights from new therapeutics. Nature Reviews Drug Discovery 21, 306–318 (2022).

3. C. Kurz, L. Walker, B. S. Rauchmann, R. Perneczky, Dysfunction of the blood-brain barrier in Alzheimer’s disease: Evidence from human studies. Neuropathol Appl Neurobiol 48, e12782 (2022).

4. Z. Arvanitakis, A. W. Capuano, S. E. Leurgans, D. A. Bennett, J. A. Schneider, Relation of cerebral vessel disease to Alzheimer’s disease dementia and cognitive function in elderly people: a cross-sectional study. Lancet Neurol 15, 934–943 (2016).

5. K. Kisler, A. R. Nelson, A. Montagne, B. V. Zlokovic, Cerebral blood flow regulation and neurovascular dysfunction in Alzheimer disease. Nat Rev Neurosci 18, 419–434 (2017).

6. Y. Iturria-Medina et al., Early role of vascular dysregulation on late-onset Alzheimer’s disease based on multifactorial data-driven analysis. Nat Commun 7, 11934 (2016).

7. J. M. Wardlaw et al., Neuroimaging standards for research into small vessel disease and its contribution to ageing and neurodegeneration. Lancet Neurol 12, 822–838 (2013).

8. K. A. Jellinger, Alzheimer disease and cerebrovascular pathology: an update. J Neural Transm (Vienna) 109, 813–836 (2002).

9. R. E. Bennett et al., Tau induces blood vessel abnormalities and angiogenesis-related gene expression in P301L transgenic mice and human Alzheimer’s disease. Proc Natl Acad Sci U S A 115, E1289–E1298 (2018).

10. Y. Moon et al., Blood-brain barrier breakdown is linked to tau pathology and neuronal injury in a differential manner according to amyloid deposition. J Cereb Blood Flow Metab 43, 1813–1825 (2023).

11. R. A. Fisher, J. S. Miners, S. Love, Pathological changes within the cerebral vasculature in Alzheimer’s disease: New perspectives. Brain Pathol 32, e13061 (2022).

12. D. A. Nation et al., Blood-brain barrier breakdown is an early biomarker of human cognitive dysfunction. Nat Med 25, 270–276 (2019).

13. M. D. Sweeney, K. Kisler, A. Montagne, A. W. Toga, B. V. Zlokovic, The role of brain vasculature in neurodegenerative disorders. Nat Neurosci 21, 1318–1331 (2018).

14. Q. Liu et al., Amyloid precursor protein regulates brain apolipoprotein E and cholesterol metabolism through lipoprotein receptor LRP1. Neuron 56, 66–78 (2007).

15. T, Kanekiyo, & G, Bu The low-density lipoprotein receptor-related protein 1 and amyloid-β clearance in Alzheimers disease. Front. Aging Neurosci. 6,93(2014) and tau

16. J. N. Rauch, G. Luna, E. Guzman et al. LRP1 is a master regulator of tau uptake and spread. Nature 580, 381–385 (2020).

17. J. M. Cooper et al., Regulation of tau internalization, degradation, and seeding by LRP1 reveals multiple pathways for tau catabolism. J Biol Chem 296, 100715 (2021).

18. R. Deane et al. LRP/Amyloid β-Peptide interaction mediates differential brain efflux of Aβ isoforms. Neuron 43, 333–344 (2004).

19. K. Yamada et al. The low density lipoprotein receptor-related protein 1 mediates uptake of amyloid β peptides in an in vitro model of the blood-brain barrier cells. J. Biol. Chem. 283, 34554–34562 (2008).

20. L. B. Jaeger et al. Testing the neurovascular hypothesis of Alzheimer’s disease: LRP-1 antisense reduces blood-brain barrier clearance, increases brain levels of amyloid-β protein, and impairs cognition. J. Alzheimer’sDis. 17,553–570 (2009);

21. T. Kanekiyo et al. Neuronal clearance of amyloid-β by endocytic receptor LRP1. J. Neurosci. 33,19276–19283 (2013).,

22. S. E. Storck et al. Endothelial LRP1 transports amyloid-β1-42 across the blood-brain barrier. J. Clin. Investig. 126,123–136 (2016)., 5,15–21

23. L. B. Jaeger et al. Testing the neurovascular hypothesis of Alzheimer’s disease: LRP-1 antisense reduces blood-brain barrier clearance, increases brain levels of amyloid-β protein, and impairs cognition. J. Alzheimer’sDis. 17,553–570 (2009);

24. A. Montagne, et al. Blood-brain barrier breakdown in the aging human hippocampus. Neuron 85,296–302 (2015);

25. A. Montagne et al. APOE4 leads to blood-brain barrier dysfunction predicting cognitive decline. Nature 581,71–76 (2020);

26. A. M. Nikolakopoulou et al. Endothelial LRP1 protects against neurodegeneration by blocking cyclophilin A. J. Exp. Med. 218,11–21 (2021);

27. B. Van Gool et al. LRP1 has a predominant role in production over clearance of Aβ in a mouse model of Alzheimer’sdisease. Mol. Neurobiol. 56,7234–7245 (2019),

28. K. Govindpani, L. G. McNamara, N. R Smith, C. Vinnakota, H. J. Waldvogel, R. L.M. Faull, and A. Kwakowsky, 2019. “Vascular dysfunction in alzheimer’s disease: a prelude to the pathological process or a consequence of it?”, Journal of Clinical Medicine, 8:651 (2019)

29. Y Tian, J. C. Chang, E. Y. Fan, M. Flajolet, & P. Greengard, P. Adaptor complex AP2/ PICALM, through interaction with LC3, targets Alzheimer’s APP-CTF for terminal degradation via autop-hagy. Proc.NatlAcad.Sci. 110,17071–17076 (2013);

30. Z, Zhao, et al. Central role for PICALM in amyloid-β blood-brain barrier transcytosis and clearance. Nat. Neurosci. 18, 978–987 (2015).

31. S. Storck, et al. “The concerted amyloid-beta clearance of lrp1 and abcb1/p-gp across the blood-brain barrier is linked by picalm”. Brain Behavior and Immunity, vol. 73, p. 21–33. (2018).

32. X. Tian et al., On the shuttling across the blood-brain barrier via tubule formation: Mechanism and cargo avidity bias. Sci Adv 6, (2020).

33. M. L. D et al., Syndapin-2 mediated transcytosis of amyloid-beta across the blood-brain barrier. Brain Commun 4, fcac039 (2022).

34. X. Tian, S. Nyberg, P. Sharp, et al. LRP-1-mediated intracellular antibody delivery to the Central Nervous System. Sci Rep 5, 11990 (2015)

35. X. H. Tian, S. Angioletti-Uberti, G. Battaglia, On the design of precision nanomedicines. Science Advances 6, (2020).

36. C. H. van Dyck et al., Lecanemab in Early Alzheimer’s Disease. N Engl J Med 388, 9–21 (2023).

37. N. J. Reish et al., Multiple Cerebral Hemorrhages in a Patient Receiving Lecanemab and Treated with t-PA for Stroke. N Engl J Med 388, 478–479 (2023).

38. S. Reardon, Alzheimers drug donanemab what promising trial means for treatments. 617, (2023).

39. C. M. Clark et al., Use of Florbetapir-PET for Imaging β-Amyloid Pathology. Jama-Journal of the American Medical Association 305, 275–283 (2011).

