## supporting information for "Multivalent modulation of endothelial LRP1 induces fast neurovascular amyloid-β clearance and cognitive function improvement in Alzheimer’s disease models"

### Multivalent Targeting of Blood-Brain Barrier LRP1 for Neurovascular Recovery Therapy for Alzheimer's Disease

*[10]New address: Curapath, Av. Benjamín Franklin, 19, 46980 Paterna, Valencia*

### Supporting information

#### 1. Materials and methods

##### $A_{39}$-POs Polymersome Preparation and Characterisation

$P[(OEG)_{10}MA]_{20}$-$PDPA_{120}$ and Angiopep2-$P[(OEG)_{10}MA]_{20}$-$PDPA_{120}$ copolymers were synthesised via atom transfer radical polymerisation (ATRP) as previously reported (1). $A_{39}$-POs were formulated utilising the film-rehydration method, whereby $P[(OEG)_{10}MA]_{20}$-$PDPA_{120}$ and a 1.88 % molar ratio of Angiopep2-$P[(OEG)_{10}MA]_{20}$-$PDPA_{120}$ were dissolved in a methanol and chloroform mixture (v/v, 3 : 1). The mixture was left to evaporate, allowing the organic solvent to completely dissipate and form a homogeneous polymer film at the bottom of the vessel. PBS solution was added to rehydrate the polymer film (10 g/L) and sonicated for 30 min. The solution was stirred continuously for 7 days at 4°C using a magnetic stirrer at 1000 rpm. The morphology of $A_{39}$-POs was examined via transmission electron microscopy (JEM-2100Plus, Japan). Diameter distribution was assessed by dynamic light scattering (Malvern Zetasizer Pro, UK).

##### Animals

All animal studies were conducted in accordance with the guidelines set forth by the *West China Hospital Animal Care Committee*. Considerable efforts were made to minimise the number of animals utilised in these studies and to alleviate any pain or discomfort experienced by the animals. In all experiments, animals were accommodated in a room where the temperature was regulated, with consistent alternating cycles of light and darkness.

**Immunohistochemistry (IHC)**

Paraffin-embedded sections (5 μm) of mouse brain tissue were deparaffinised with xylene and subjected to gradient alcohol hydration (30%, 50%, 80%, 100%). Endogenous peroxidase activity was quenched using 3% $H_2O_2$ (room temperature, 10 min) following antigen retrieval (Biosharp, 22315828). The tissues were blocked with 1% BSA (Aladdin, A104912) at room temperature for 30 min. Aβ targeting primary antibodies were applied overnight at 4°C (Servicebio, GB13414-1, 1:200), followed by washing with PBS. The sections were then incubated with HRP-conjugated secondary antibodies (Abclonal, AS014, 1:150) at ambient temperature for 30 min and subsequently washed with PBS. DAB substrate (Elabscience, E-IR-R101, 1:20) and haematoxylin differentiation solution (Servicebio, G1004) were sequentially applied to the sections for 5 min each. The sections were further treated with haematoxylin solution (Servicebio, G1039) at room temperature for 1 min, followed by a PBS wash. After another PBS wash, sections were counterstained with Dako bluing buffer (Servicebio, G1040) at room temperature for 1 min. Lastly, the slides were mounted using a quick-drying neutral resin (ZSZSGBBIO, ZLI-9516).

**Aβ Extraction**

Following euthanasia via an overdose of isoflurane anaesthesia, cardiac perfusion was conducted with cold PBS. Subsequently, the entire brain tissue was harvested and weighed. The brain tissues underwent comprehensive homogenisation (Servicebio, KZ-5F-3D) employing a TBS solution imbued with phosphatase inhibitor (Servicebio, CR2302054) and protease inhibitor (Servicebio, CR2306008) at 70 Hz, -20°C. The supernatant was segregated subsequent to centrifuging the brain tissue homogenate for 1 h at 100,000g and 4°C utilising an ultra-high-speed centrifuge (Beckman, Optima MAX-XP). The tissue residue was then reprocessed. The residues were resuspended in 70% formic acid (Aopusheng (Tianjin) Chemical, 20210610), centrifuged at 100,000g, 4°C for 1 h, and the resultant supernatant was collected.

**Vessel Extraction**

Immediately following euthanasia via isoflurane overdose anaesthesia, the heart was perfused with cold PBS, and brain tissues were excised and weighed. The tissues were processed using a vascular parenchyma isolation buffer comprising 10 mM HEPES (Servicebio, CR2207064), 141 mM NaCl (Aladdin, 111549), 4 mM KCl (Aladdin, P112134), 2.8 mM $CaCl_2$ (Aladdin, C290953), 1 mM $MgSO_4$ (Aladdin, M433513), 1 mM $NaH_2PO_4$ (Aladdin, S433623), and 10 mM glucose (Servicebio, CR2112094). Fresh brain tissue underwent vascular and parenchymal separation. To 500 μL of brain tissue homogenate, 1 mL of 26% (w/w) dextran (Next Sage, 61212ES60) was added. Following thorough mixing and a resting period of 15 min at 4°C, the mixture was centrifuged (15,800g, 15 min, 4°C). Subsequently, the upper layer represented the parenchymal fraction, and the lower layer corresponded to the vascular fraction, with both fractions being thoroughly rinsed with PBS.

**Enzyme-Linked Immunosorbent Assay (ELISA)**

The already extracted brain parenchyma and blood vessels were dissolved at room temperature. The parenchyma and blood vessels underwent comprehensive homogenisation using a tissue lysis solution (Yase, 016c1050) enhanced with a phosphatase inhibitor (Servicebio, CR2302054) and a protease inhibitor (Servicebio, CR2306008) at 70 Hz, -20°C. The tissues were allowed to fully lyse by resting at 4°C for 15 min. The supernatant was collected via centrifugation at 14,000 g, 4°C for 10 min. Protein concentrations for various proteins, including LRP1 (NOVUS, NBP3-00449), PACSIN2 (Anruike,

YX-160103M), RAB5 (abbexa, abx154598), GLUT1 (Anruike, YX-071242M), Claudin11 (Anruike, YX-031229M), ZO1 (Elabscience, E-EL-M1161), MBP (Elabscience, E-EL-M0805), MAP-2 (Anruike, YX-120118M), SRB1 (Anruike, ARK-756321A), and Aβ (invitrogen, KMB3441) were determined using ELISA kits following the manufacturer protocols. For Aβ in blood the test was performed directly according to the ELISA instructions. Aβ extracted by formic acid needs to be neutralized with tris base before test. Protein concentrations were calculated based on the curve equation (four-parameter fit).

**Confocal Imaging**

Confocal images were captured using a Leica Stellaris 5 confocal microscope, equipped with Diode 405, Argon, DPSS 561, and HeNe633 lasers. Imaging was conducted at a resolution of 2048×2048 pixels and a scanning speed of ×100. Colocalisation analysis to derive Pearson's correlation coefficient r was performed utilising the colocalisation plug-in on ImageJ.

**Tyramide Signal Amplification (TSA) Stain**

Mouse brain tissue sections were deparaffinised with xylene and hydrated through gradient alcohol dehydration. Following antigen retrieval, 3 % $H_2O_2$ was utilised to inactivate endogenous peroxidase in tissues (Biosarp,

22315828) for 10 min. Permeabilisation of sections at room temperature for 10 min was achieved using 1% Triton X-100 (Biofroxx, 1139ML100). The sections were rinsed with PBS and incubated with the primary antibody solution for 1 h at room temperature. After washing with PBS, the samples were incubated with the HRP-conjugated secondary antibody for 10 min. Subsequent to a PBS rinse, the TSA fluorescent solution (Absin, abs50031) was applied and incubated at room temperature for 10 min. Antigen repair was subsequently conducted. The aforementioned steps were reiterated with distinct primary antibodies, including LRP1 (abclonal, A1439, 1:100), Aβ (servicebio, GB13414-1, 1:200), CD31 (cellsignaling, 77699, 1:300), RAB5 (thermofisher, PA5-88260, 1:200) and CD146 (abcam, ab75769, 1:300), and various wavelengths of TSA, encompassing TSA520, TSA570, TSA620, and TSA700 dyes until multimeric fluorescent staining was achieved. Upon completion of the final staining cycle, nuclei were stained with DAPI (Solarbio, C0065) for 5 min, then sealed with antifade sealing agent (Solarbio, S2100).

**Positron Emission Tomography-Computed Tomography (PET-CT) Imaging**

APP/PS1 POs group (12-month-old) mice were injected intravenously with commercial Aβ radiocontrast agent [$^{18}$F]AV-45 (2.8-3.2 MBq). One hour later, intracranial images of the mice were acquired using a Micro-PET-CT imager

(Inviscan, France). After a recovery period of 3 weeks, saline (200 μL) was intravenously injected to wild-type mice (3- and 12-months-old) and sham APP/PS1 mice (12-months-old). Meanwile APP/PS1 POs group were intravenously injected 200 μL of $A_{39}$-POs (10 g/L). 12 hours later, all mice were injected intravenously with [$^{18}$F]AV-45 (2.8-3.2 MBq). One hour later, intracranial images of the mice were acquired using the same imager.

**Hematoxylin-Eosin (H&E) Staining**

Mouse tissue sections (5 μm) were dewaxed and rehydrated. The sections were stained with haematoxylin solution for 5 min, followed by incubation with differentiation solution (Servicebio, G1039) for 10 - 15 s. Subsequently, the sections were treated with Dako bluing solution (Servicebio, G1040) for 30 s and washed with PBS. After air-drying, the sections were stained with eosin staining solution for 30 s, followed by washing with ethanol. The slides underwent dehydration through a gradient of ethanol (30%, 50%, 80%, 100%) and xylene. Finally, the tissue sections were sealed with a quick-drying neutral resin (ZSZSGBBIO, ZLI-9516).

##### Tissue Clearing and Staining

The paraformaldehyde-fixed mouse brain tissues were treated with 1/2 CUBIC-L solution (80 mL Milli-Q, 5 g Triton X-100, 5 g N-butyldiethanolamine (Aladdin, 102-79-4)) at 37°C for 6 h. Then, the solution was replaced with a CUBIC-L solution and continued to be treated at 37°C for 15 days, with the new CUBIC-L solution being replaced every 2 days. Then the brain tissues were washed three times with staining buffer (1.5 M NaCl). The hyalinised brains were transferred to a staining buffer containing vascular probe (Vector, DL-1178-1, 1:100) and Aβ probe (abcam, ab216983, 100 nM) for fluorescence staining (RT, 3 days). The refractive index was adjusted using CUBIC-M solution (25 g Milli-Q, 45 g antipyrine (Aladdin, 60-80-0), 30 g N-methylnicotinamide (Aladdin, 114-33-0), and 125 μl of N-butyldiethanolamine)). Brain tissues were stored in dibenzyl ether (Sigma-Aldrich) until light sheet imaging. Images were analysed using Amira and iMaris software.

##### High Performance Liquid Chromatography (HPLC)

Donepezil HCl (Aladdin, D129948) dissolved in PBS and added to a polymer film to prepare donepezil HCl@$A_{39}$-POs, followed by dialysis using a 3 kDa dialysis bag for 7 days. The concentration of donepezil HCl in the dialysis fluid was measured to calculate the encapsulation efficiency. Sodium 1-

decanesulfonate (Aladdin, S100284) was dissolved in pure water (15.7416 mM/L) and filtered. Chromatographic grade acetonitrile solution and perchloric acid were added, followed by ultrasonication for 10 min. Standard solutions of donepezil HCl at concentrations of 250, 125, 62.5, 31.25, 15.625, and 7.8125 μg/mL were prepared. The content of donepezil HCl was detected using an Agilent-1260 chromatograph under the conditions of a flow rate of 1 mL/min, column temperature of 35°C, volume of 20 μl, and detection signal at 271 nm.

**Morris Water Maze (MWM) Experiment**

In each group, mice were administered a caudal vein injection daily ($A_{39}$-POs, Donepezil@$A_{39}$-POs, Donepezil, or saline), continuing for three consecutive days. Subsequently, mice were housed in a standard rearing environment for seven days to acclimatise and recover. For the analysis, the pool was segmented into four quadrants. From days 11 to 14 (days 375th - 378th of lifespan), a platform was positioned in the II quadrant, and animals were introduced into the thermostatic pool from the midpoint of each quadrant daily, with time taken by the mice to locate the platform recorded as escape latency. Should mice fail to reach the platform within 120 s, they were guided to it and remained there for 30 s. Spatial probe (Stage II) was conducted on day 15 (day 379th of lifespan). With platform removed, mice were placed into water from the midpoint of IV quadrant. The results of spatial probe were expressed as either the percentage of time the mice remained at the original escape platform location or the number of times they passed. Reverse Place Navigation trials were conducted from days 16 to 19 (day 380th - 383th of lifespan), with the platform positioned in the quadrant opposite the original platform location (IV quadrant). The animals repeated the regimen from days 11 to 14, and reverse escape latency was documented. On day 20 (day 384th of lifespan), the platform was removed, and mice were introduced into the water from the midpoint of II quadrant. The results of  Reverse spatial probe were expressed as either the percentage of time that the animal spent on the escape platform or the number of times it crossed the original platform position. In addition, six months later the mice were subjected to place navigation (Stage V) and spatial probe (Stage VI) experiments again with same methods and conditions. Animal performances were recorded by same tracking system (Ethovision XT, Noldus Information Technology) for each stage.

**Sucrose preference**

At the end of Stage IV and stage VI Sham APP/PS1 and APP/PS1 POs mice were tested for sucrose preference. Each mice were housed individually and allowed to acclimatize the cage containing two bottles of standard pure water for 2 days. One of the bottles of standard purified water was subsequently replaced with 2% sucrose solution. The amount of water consumed by mice was recorded and refresh water for each bottle daily. Sucrose preference = (Sucrose water consumption / Sucrose water consumption + Standard purified water consumption) x 100%

**Nest construction**

3 days after Sucrose preference experiment, the mice Each mice were housed individually and then adapted to a single cage environment for 7 days. Subsequently, the 10 pieces of paper ($5 \times 5$ $cm^2$) were added in each cage and evenly placed. After 4 days, the nesting test was scored in line with the improved 4-point system ranging (2). 1 point, no visible tear, no recognizable nest site; 2 points, no visible tear, nest site recognizable; 3 points, partial tear, recognizable nest site; 4 points, sharpest tear, recognizable nest.

#### 2. Extended figures

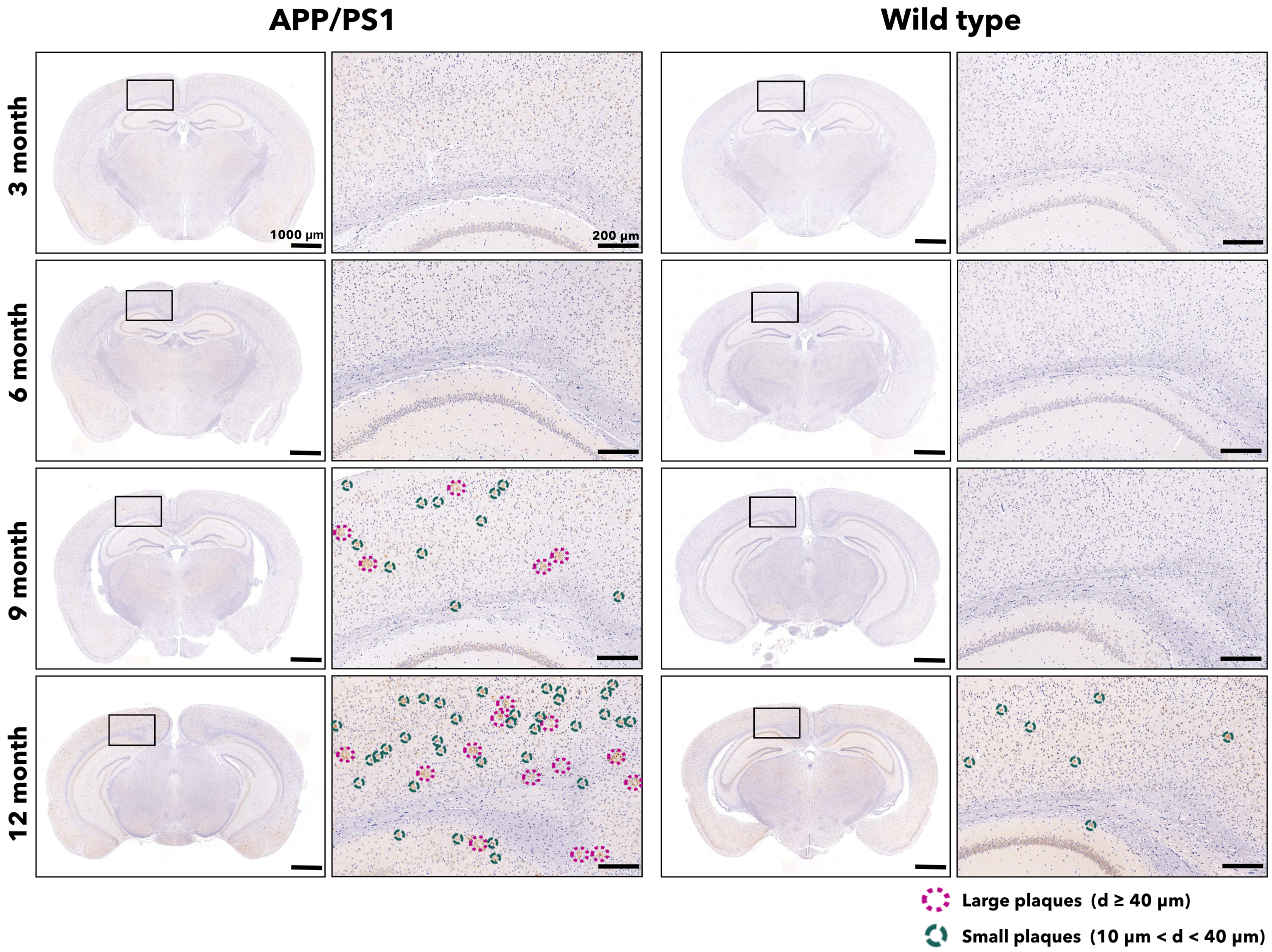

**Fig. S1. IHC images of Aβ in the brains sections of age-matched mice.**

IHC images of Aβ in coronal brain sections from mice at 3, 6, 9, and 12 months old. Each section is derived from a different mouse brain. As age increases, area percentage of the brain occupied by Aβ increases in both APP/PS1 and wild type mice (n = 3).

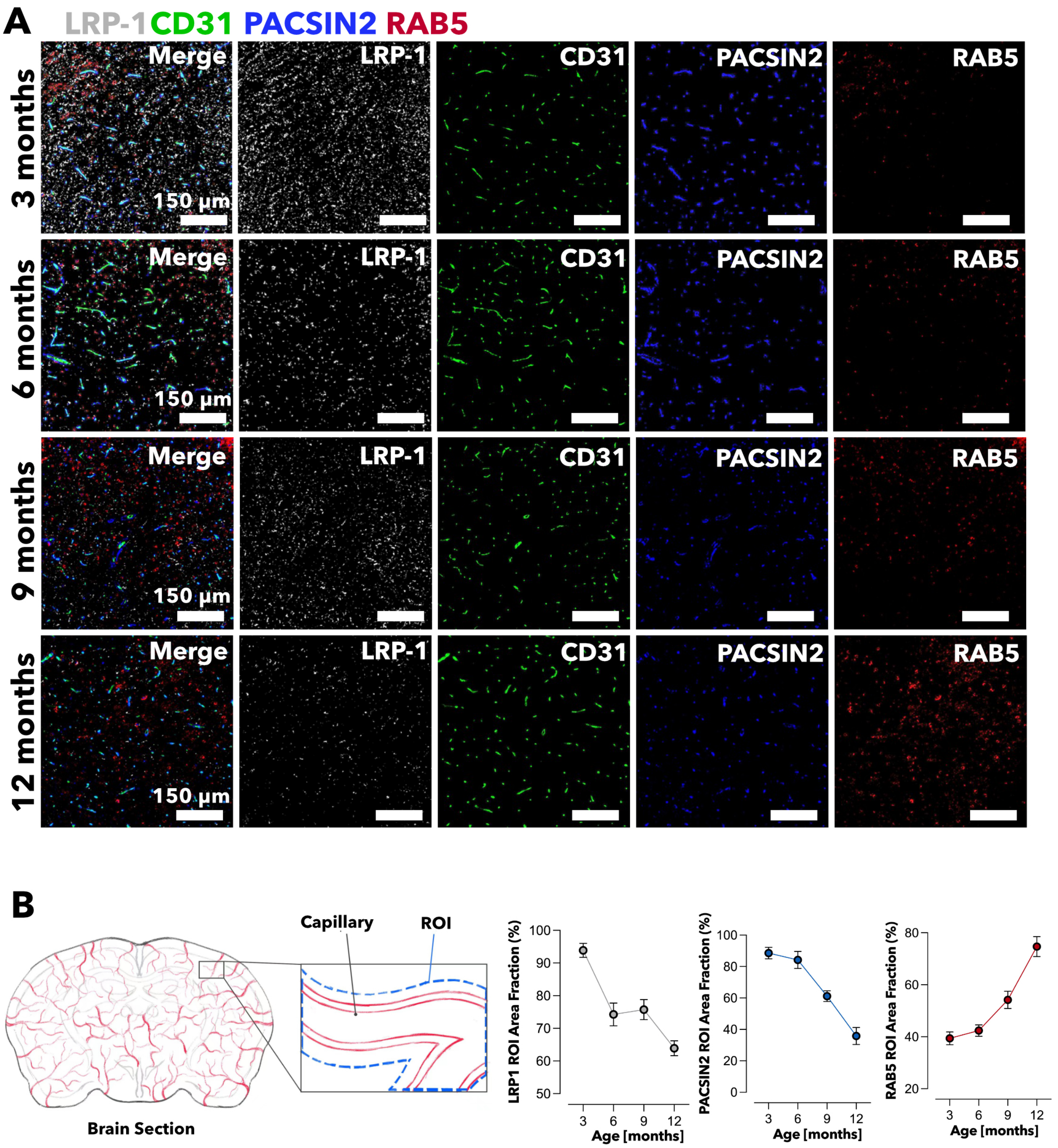

**Fig. S2. Fluorescence micrographs colocalization analysis of age-matched brain sections.**

Fluorescence images of brain sections from 3, 6, 9, and 12 months old wild-type mice, LRP1 (grey) CD31 (green) PACSIN2 (blue) RAB5 (red) (A). Capillary regions were counted as ROIs, and the percentage area of LRP1, PACSIN2, and RAB5 in the ROIs was counted. A total of 10 capillaries from two independent trials were selected for counting. LRP1, PACSIN2, and RAB5 showed the same trend as in the ELISA assay (B).

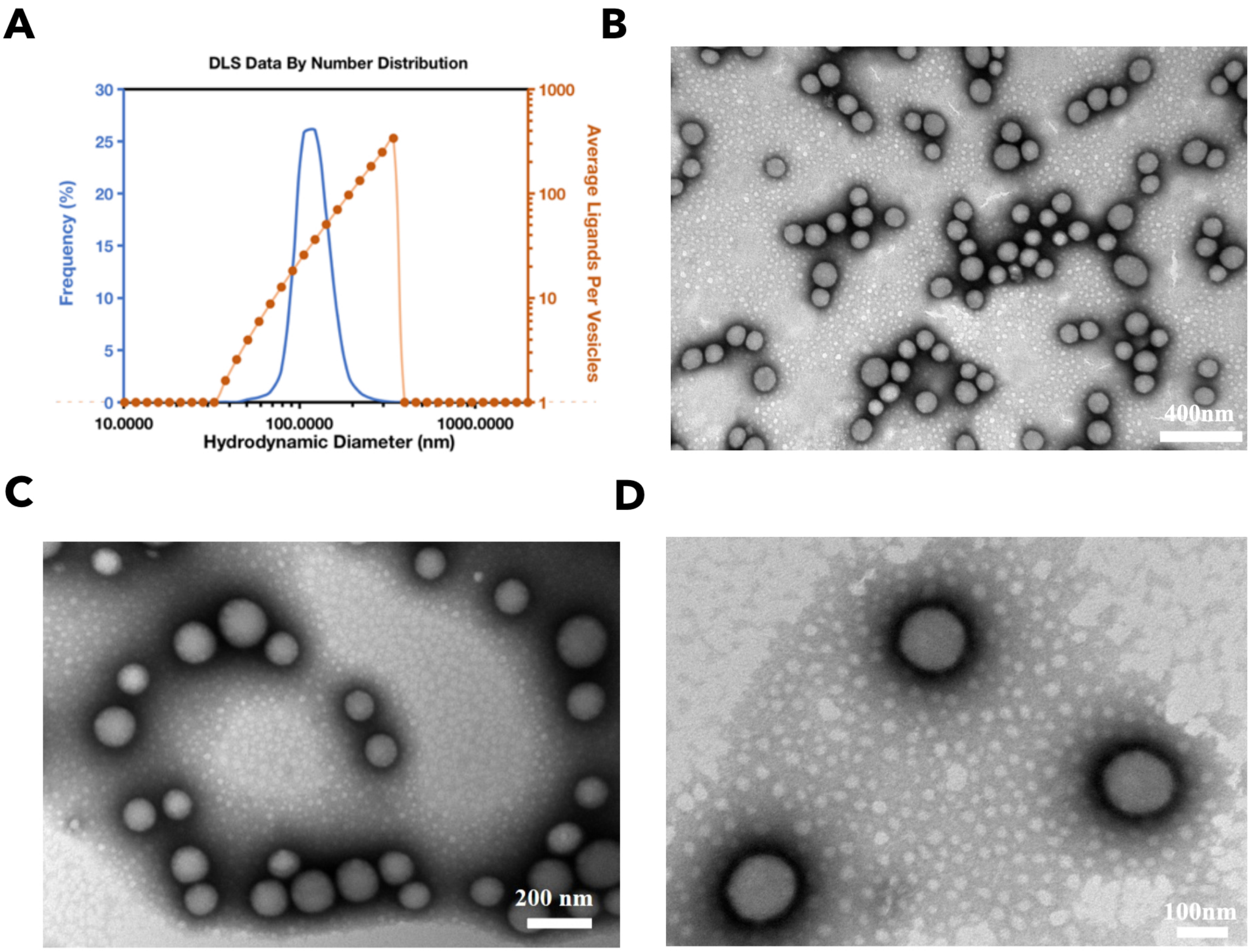

**Fig. S3. Characterization of $A_{39}$-POs in terms of size and morphology.**

Size distribution and diameter matched ligands number of $A_{39}$-POs (A). TEM images of $A_{39}$-POs magnified × 60,000 (B), × 100,000 (C) and × 140,000 (D).

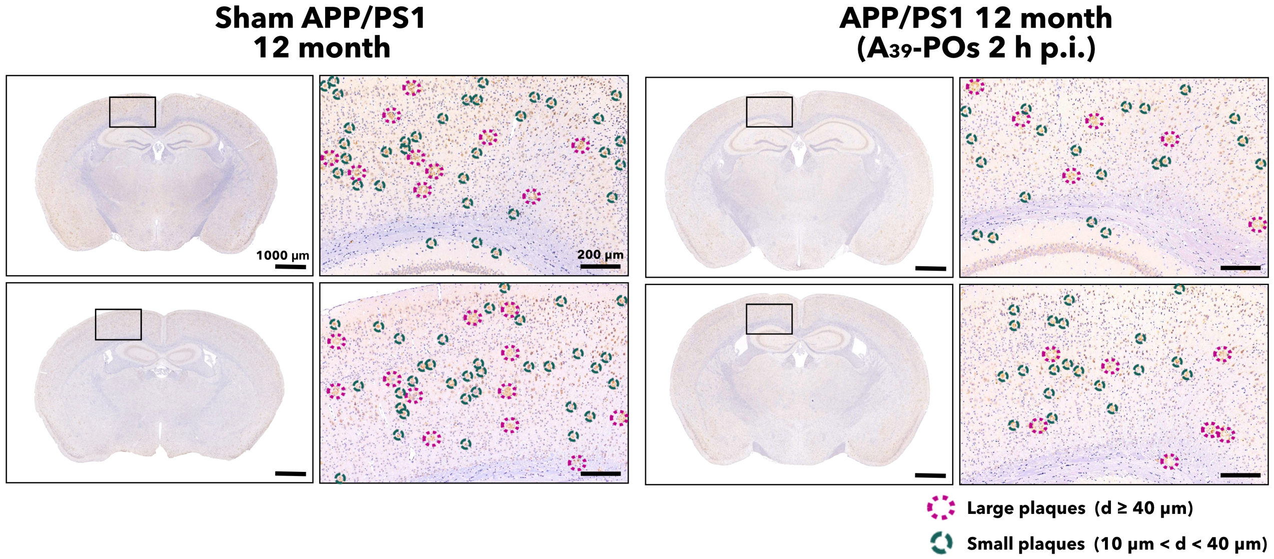

**Fig. S4. IHC images of Aβ in the brain sections of 12-month-old Sham APP/PS1 and $A_{39}$-POs treated mice.**

IHC images of Aβ in coronal brain sections from 12-month-old mice. Each section is derived from a different mouse brain. The percentage of the brain area occupied by Aβ plaques reduced after 2 hr post $A_{39}$-POs injection. Meanwhile, both the number of large and small plaques decreased.

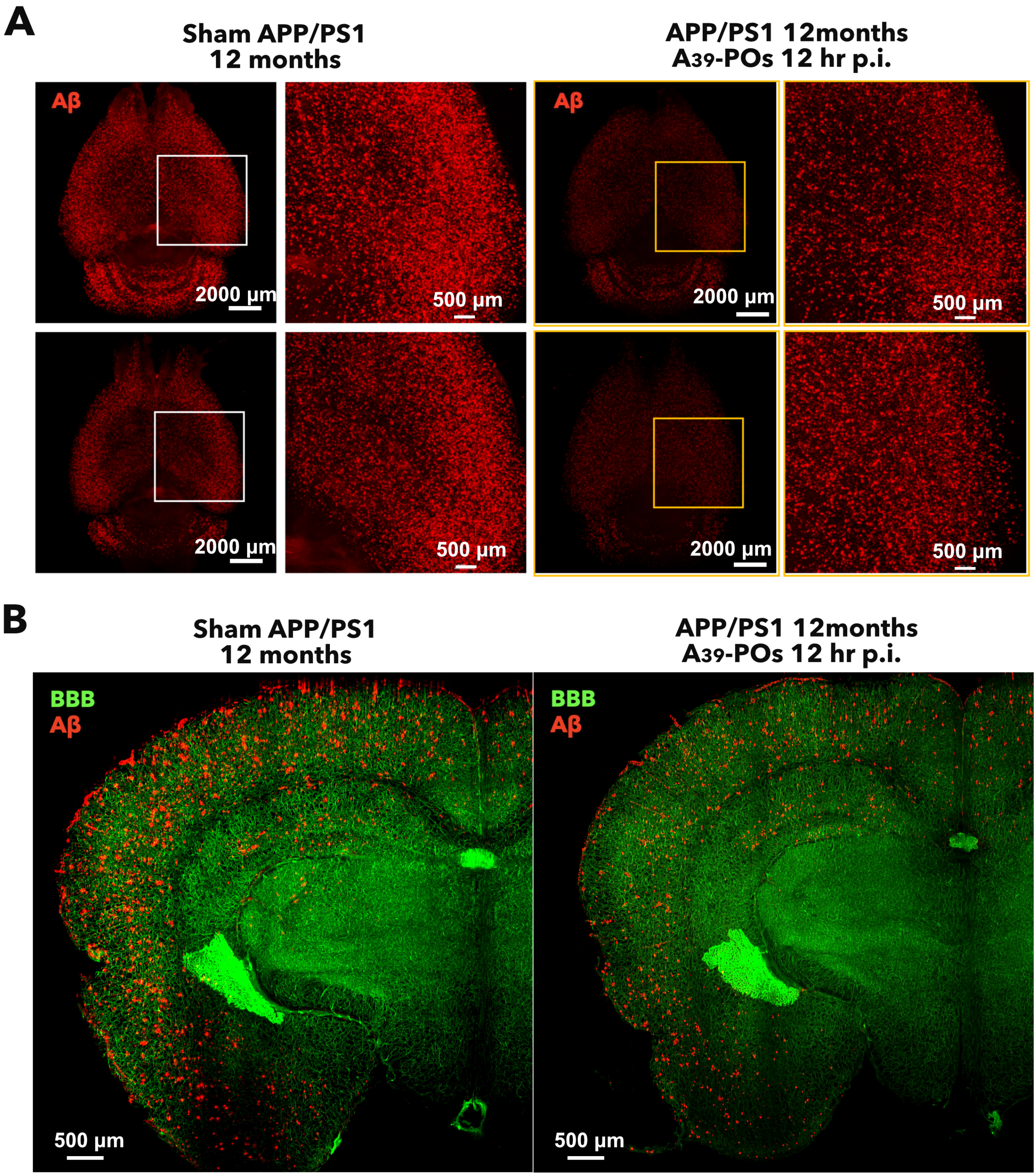

**Fig. S5. Brain clearing images of 12-month-old Sham APP/PS1 and $A_{39}$-POs treated mice.**

3D brain clearing images of Aβ (red) in the mice brain (A). Coronal view of mouse brain with a thickness of 300 μm (BBB in green, Aβ in red) (B). The Aβ signal is attenuated in the brains treated with $A_{39}$-POs.

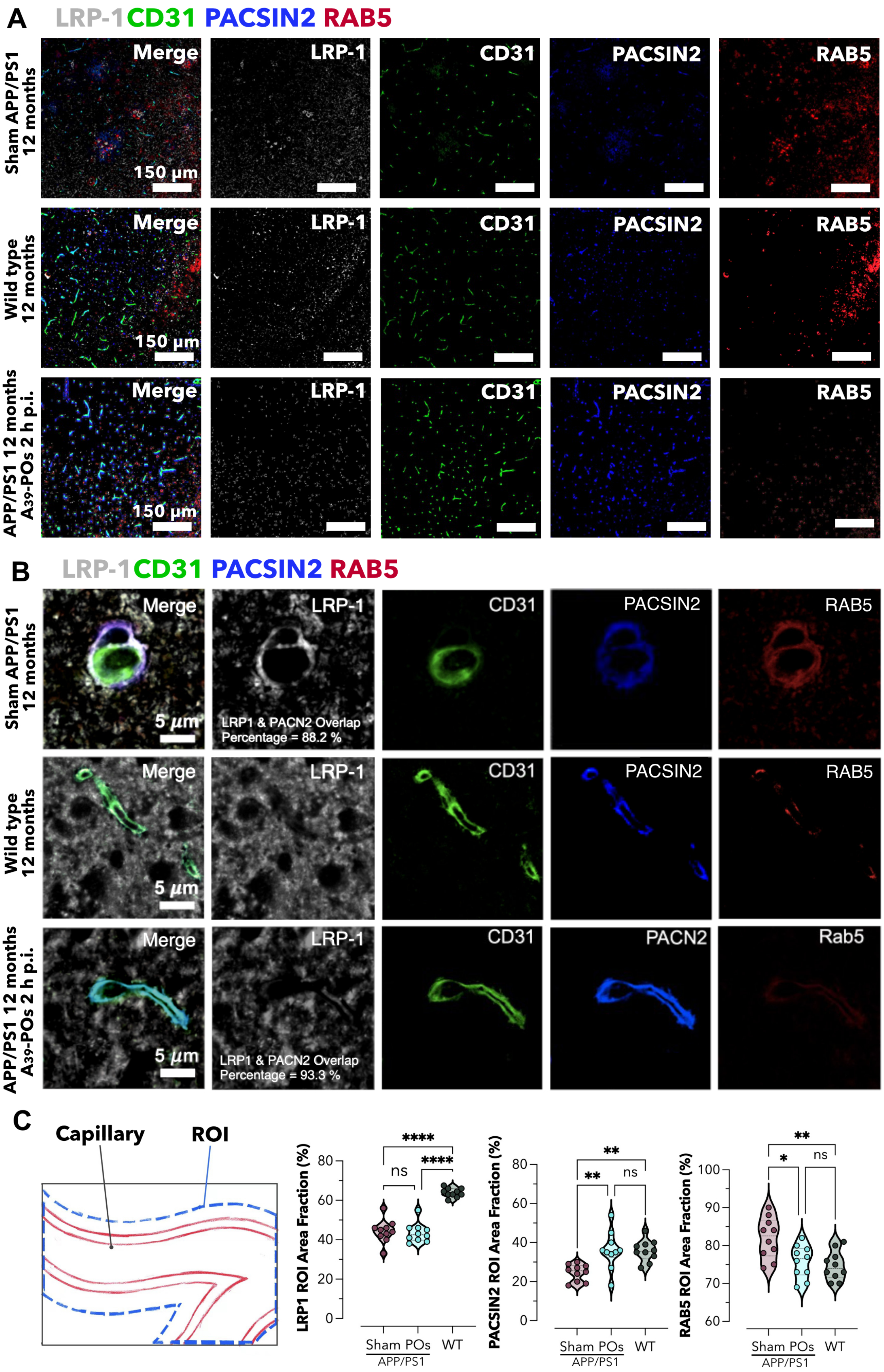
A
LRP-1 CD31 PACSIN2 RAB5
Sham APP/PS1 12 months
Wild type 12 months
APP/PS1 12 months A39-POs 2 h p.i.
Merge
LRP-1
CD31
PACSIN2
RAB5
150 µm
B
LRP-1 CD31 PACSIN2 RAB5
5 µm
LRP1 & PACN2 Overlap Percentage = 88.2 %
LRP1 & PACN2 Overlap Percentage = 93.3 %
PACN2
Rab5
C
Capillary
ROI
LRP1 ROI Area Fraction (%)
PACSIN2 ROI Area Fraction (%)
RAB5 ROI Area Fraction (%)
Sham POs WT
APP/PS1
ns
****
**
*

**Fig. S6. Fluorescence micrographs co-localisation analysis of treated brain.**

Fluorescence images of brain sections from 12-month-old wild-type AD and treated AD mice, LRP1 (grey) CD31 (green) PACSIN2 (blue) RAB5 (red) (A). Confocal microscope image of brain sections from 12-month-old wild-type AD and treated AD mice (B). Capillary regions were counted as ROIs, and the percentage area of LRP1, PACSIN2, and RAB5 in the ROIs was counted. A total of 10 capillaries from two independent trials were selected for counting. LRP1, PACSIN2, and RAB5 followed the same trend as in the ELISA assay (C).

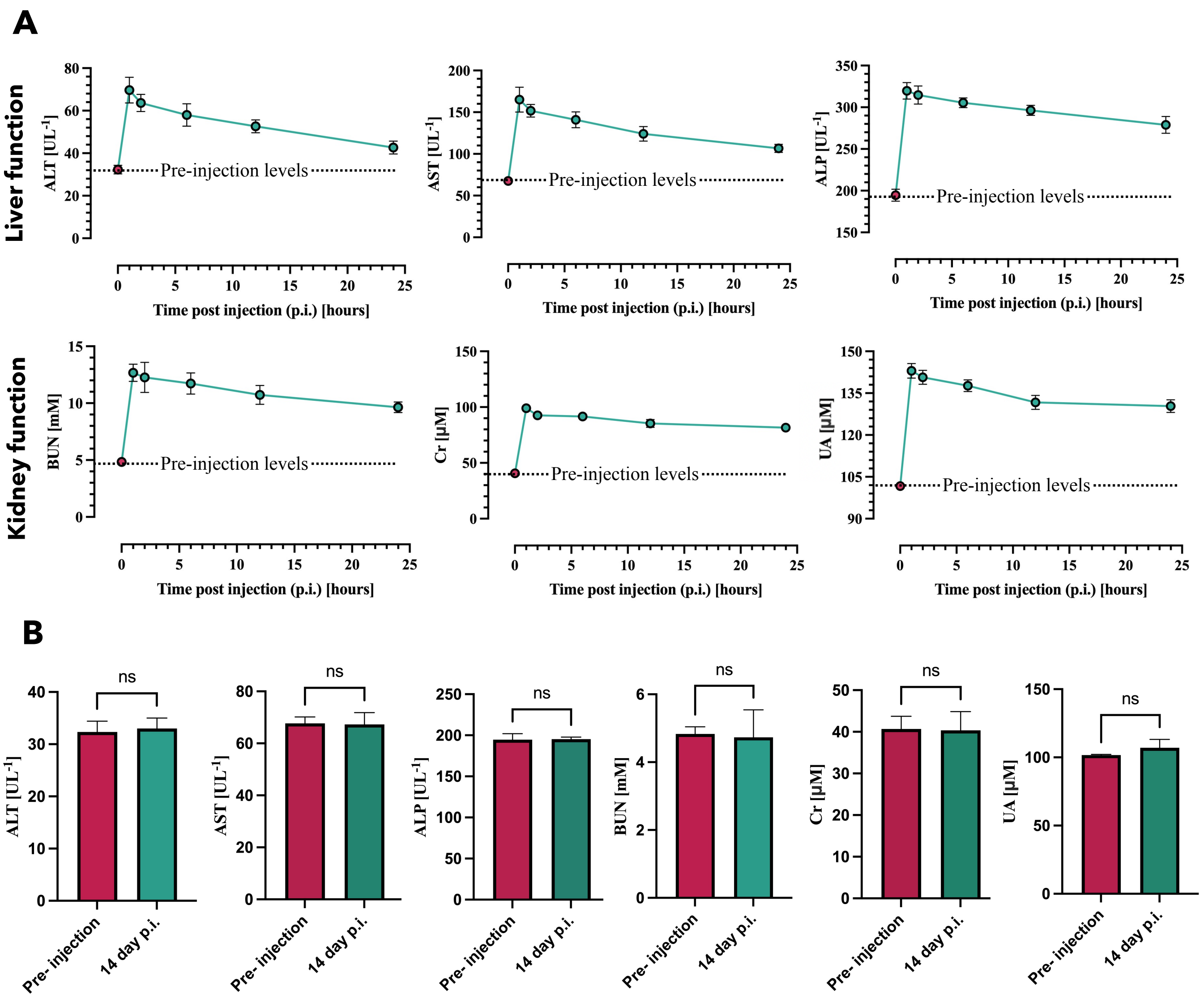

**Fig. S7. Blood biochemical tests to evaluate the biosafety of $A_{39}$-POs.**

Liver (ALT, AST and ALP) and Kidney (BUN, Cr and UA) function test of $A_{39}$-POs treated mice at time post injection of 1 hr, 2 hr, 6 hr, 12 hr and 24 hr (A) and 14 days (B).

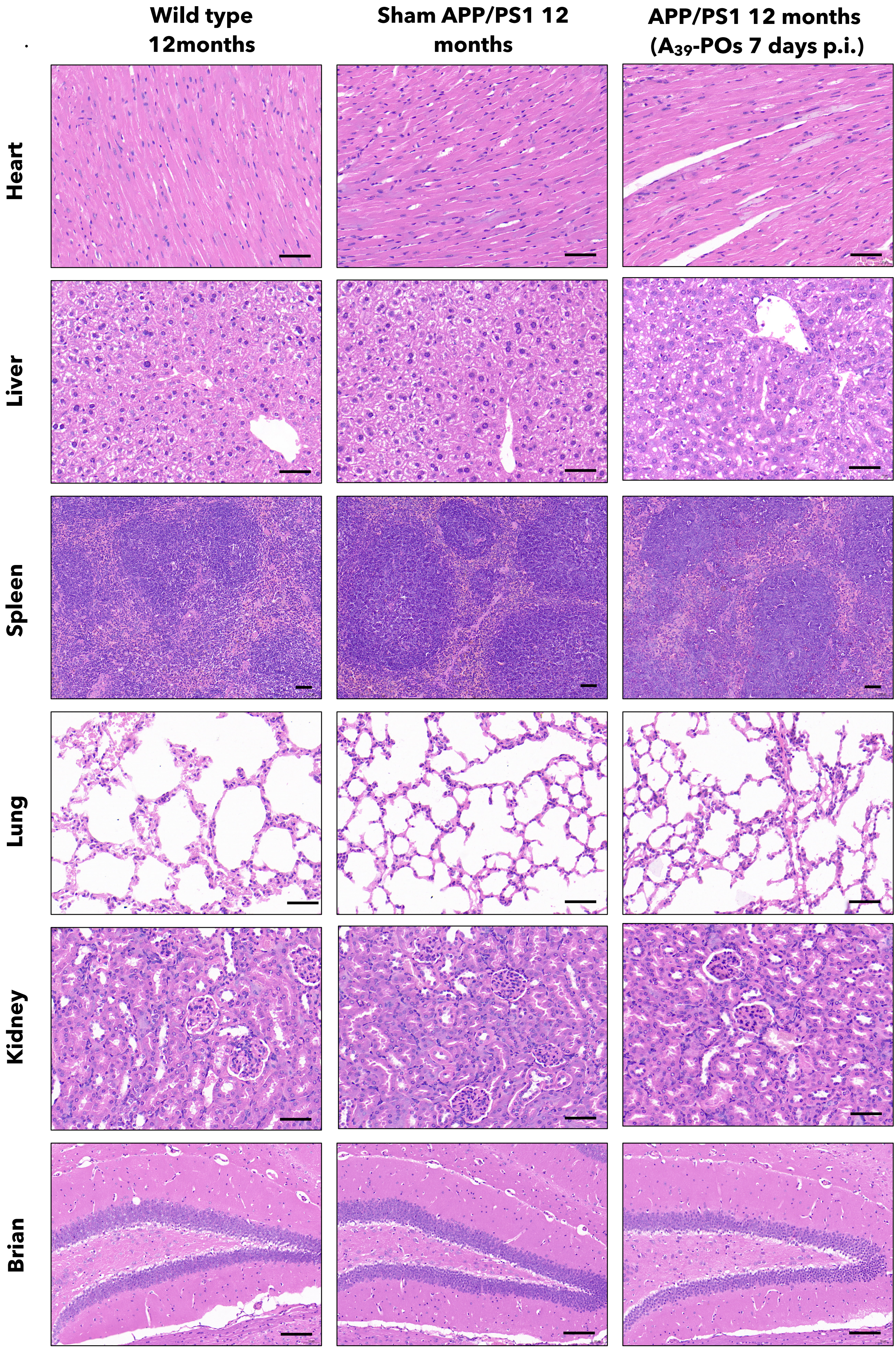

Wild type 12months
Sham APP/PS1 12 months
APP/PS1 12 months (A39-POs 7 days p.i.)
Heart
Liver
Spleen
Lung
Kidney
Brian

**Fig. S8. H&E staining to evaluate the biosafety of A39-POs.**

Heart, liver, spleen, lung, kidney, and brain H&E staining of WT, AD, and $A_{39}$-POs treated mice (7 days post injection). The cells of $A_{39}$-POs treated mice's tissues and organs did not show nuclear crumpling and cellular deformation. All Scale bars = 50 μm, animals with the age of 12 months, n = 3.

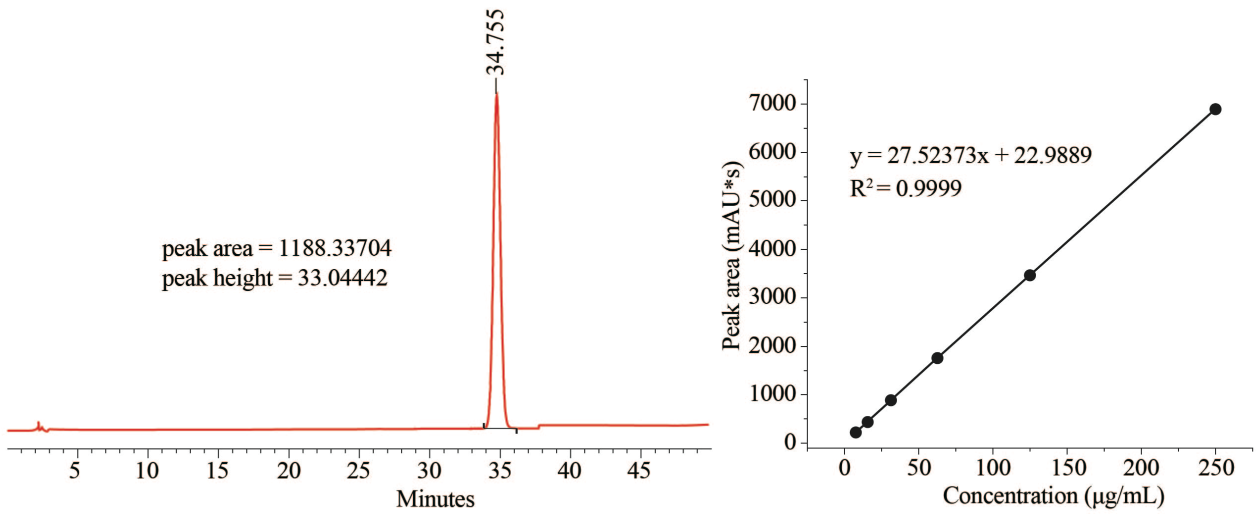

**Fig. S9. HPLC methodology was employed to ascertain the encapsulation efficiency following the encapsulation of Donepezil within $A_{39}$-POs.**

The peak area and peak height of the Donepezil@POs dialysate are depicted on the left. The corresponding standard curve is presented on the right. (Encapsulation efficiency = 20.68%)

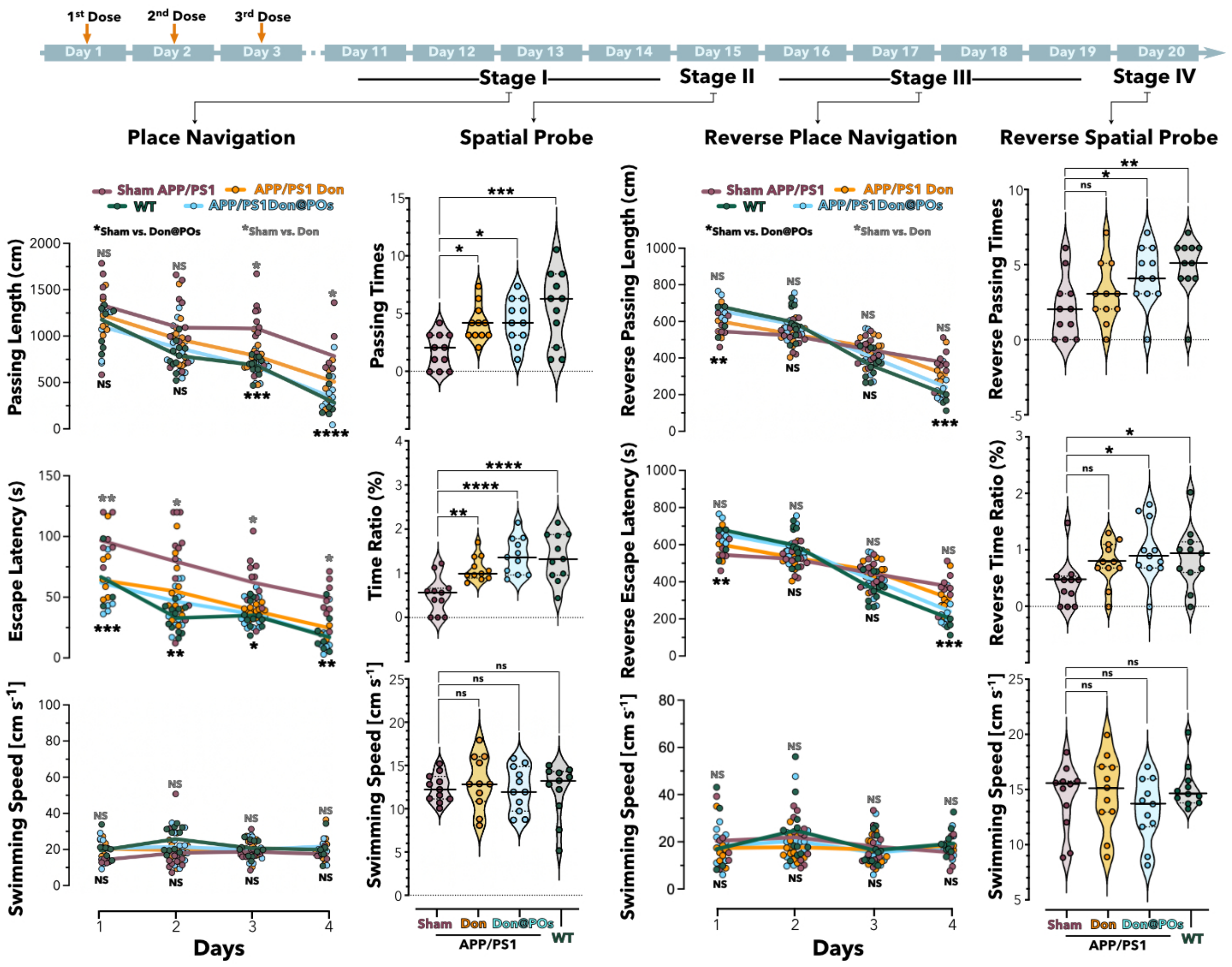

**Fig. S10. Donepezil@POs can treat AD and produce similar therapeutic effects to $A_{39}$-POs, which are superior to the same dosage of free donepezil.**

Mice were injected with saline (Sham APP/PS1 and WT group), Donepezil@POs (Don@POs group, 10g/L 200μL) or Donepezil (Don group, 2g/lL 200μL) once per day for the first three mornings. Recovery was observed from days 4 to 10 under original rearing conditions. Place navigation (Stage I) test occurred on days 11-14, showing a gradual decrease in passing length and escape latency for finding the escape platform across all groups, with the APP/PS1 Don@POs group matching the WT level and significantly outperforming the Sham APP/PS1 group. Meanwhile, the Don group demonstrated space exploration capabilities superior to Sham APP/PS1. But not to the level of the Don@POs.

Day 15th for the spatial probe (Stage II), APP/PS1 Don, APP/PS1 Don@POs, and WT groups demonstrated more passing times and a higher percentage of time spent at the escape platform's original location. The reverse place navigation (Stage III) trial from days 16-19, with the platform moved to the opposite side (IV quadrant), the APP/PS1 Don@POs and WT groups initially took longer, indicating stronger spatial memory from the stage I and stage II. However, their reverse passing length and escape latency decreased rapidly over time and were significantly lower than those of the Sham APP/PS1 mice. In this stage, which is more difficult for mice compared to stage I and II, the Don group never showed a statistically significant difference in levels from the Sham APP/PS1 group. On day 20th, in the reverse spatial probe (Stage IV) test without the platform, Don@POs treated mice still outperformed the Sham APP/PS1 group. The Don group outperformed the Sham APP/PS1 group as well, but not to the level of the WT group. Place navigation trials (Stage I and III) were analysed using two-way ANOVA, while spatial probe trials (Stage II and IV) comparisons used one-way ANOVA. Significance levels are denoted as *$p<0.05$, **$p<0.01$, ***$p<0.001$, ****$p<0.0001$, with $n \geq 11$.

#### References

1. X. Tian, S. Nyberg, P. S. Sharp, J. Madsen, N. Daneshpour, S. P. Armes, J. Berwick, M. Azzouz, P. Shaw, N. J. Abbott, G. Battaglia, LRP-1-mediated intracellular antibody delivery to the central nervous system. Sci. Rep. 5, 11990 (2015).
2. J.-W. Guo, P.-P. Guan, W.-Y. Ding, S.-L. Wang, X.-S. Huang, Z.-Y. Wang, et al., Erythrocyte membrane-encapsulated celecoxib improves the cognitive decline of Alzheimer's disease by concurrently inducing neurogenesis and reducing apoptosis in APP/PS1 transgenic mice, Biomaterials 145 (2017) 106–127.
